# Functional specialization of cytochrome *b_5_* prior to the invention of syringyl lignin biosynthesis

**DOI:** 10.1101/2023.10.18.562950

**Authors:** Xianhai Zhao, Yunjun Zhao, Qing-yin Zeng, Chang-Jun Liu

## Abstract

The production of lignin represents a hallmark of vascular plant evolution. But syringyl-lignin production is lineage specific. Its emergence in angiosperms has been regarded as a recent evolution event, as is the appearance of syringyl monomer biosynthetic F5H (CYP84A1). Angiosperm F5H uniquely requires cytochrome *b_5_* protein CB5D as an obligatory redox partner. However, it remains unclear how CB5D functionally evolved and specialized and whether it co-evolved with F5H. Here we show that plant CB5 family expanded and functionally specialized at the early stage of plant terrestrialization. CB5D was absent in green algae but emerged in bryophytes and conserved in most embryophytes, particularly proliferating in angiosperms. The innovation of CB5D is attributed to the evolution and maintenance of acidic amino residues in the helix 5 of CB5 protein. Interestingly, the syringyl-lignin producing lycophyte *Selaginella* lacks the functionally specialized CB5D. The *Selaginella* F5H (CYP788A1) solely relies on cytochrome P450 reductase as a redox partner. These results suggest that CB5 functionally diversified and specialized prior to the divergence of land plant lineages but lost in *Selaginella*; and the recently evolved angiosperm F5H has co-opted the anciently invented CB5D to form a modern cytochrome P450 monooxygenase system enabling the re-emergence of syringyl-lignin biosynthesis in higher plants.

**One Sentence Summary:** Cytochrome *b_5_* D, an indispensable electron donor for F5H-catalyzed S-lignin biosynthesis, was invented prior to the divergence of land plant lineages, earlier than the emergence of vascular plant F5Hs.

## Introduction

When the pioneering ancestors of land plants transitioned from water to terrestrial habitats approximately 470 to 600 million years ago, they immediately confronted a series of life-threatening challenges, such as exposure to damaging UV-B radiation, desiccation, temperature fluctuations, co-evolving herbivores and pathogens, and lack of structural support.(RAVEN 1984) To survive these harsh terrestrial conditions, early land plants evolved various protective mechanisms and specialized metabolic capacities. These adaptations enabled them to produce essential defense substances that facilitated their adaptation to the terrestrial ecosystem. The development of lignified vascular tissue marked a significant milestone in the evolution of land plants as they adapted to the terrestrial environment (RAVEN 1984). The deposition of lignin provided early vascular plants with physical rigidity and strengthened water-conducting cells, allowing for long-distance water transport. This innovation allowed them to significantly increase in size, ultimately contributing to their dominance in Earth’s flora (Friedman and Cook 2000; Weng et al. 2010).

Lignin is a heteropolymer primarily derived from oxidative coupling of three phenylpropanoid units, i.e., *p*-coumaryl alcohol, coniferyl alcohol and sinapyl alcohol (collectively called monolignols), which give rise to *p*-hydroxyphenyl (H), guaiacyl (G) and syringyl (S) lignin subunits when incorporated into the lignin polymer (Boerjan et al. 2003). The amount and composition of lignin monomers vary substantially among major taxa of vascular plants. In general, ferns and gymnosperms deposit lignin primarily composed of G monomers, with a small proportion of H units. In contrast, angiosperm lignins additionally contain S units, resulting in a G and S copolymers, along with some H monomers (Boerjan et al. 2003; Vanholme et al. 2010). The invention of S-lignin is regarded as a significant and characteristic evolutionary event of angiosperms (Weng et al. 2010). Interestingly, S-lignin has also been detected in *Selaginella*, a genus that represents an extant lineage of the most basal vascular plants, the lycophytes. Its emergence in *Selaginella* has been demonstrated as an independent evolutionary event as to its angiosperm counterpart (Weng et al. 2008a). Therefore, S-lignin might have evolved multiple times through convergent evolution in different plant lineages (Weng et al. 2008b; Weng et al. 2010).

In angiosperms, three cytochrome P450 monooxygenases (P450s) catalyze the hydroxylation of the phenyl ring of phenylpropanoid units, resulting in the structurally characteristic monolignols. Cinnamate 4-hydroxylase (C4H; CYP73A5) performs the *para*-hydroxylation of cinnamic acid, the initial aromatic intermediate of the phenylpropanoid pathway, producing *p*-coumaric acid. This compound is essential for the formation of all three types of lignin subunits. *p*-Coumaroyl ester 3’-hydroxylase (C3’H; CYP98A3) and ferulate/coniferyl alcohol/coniferaldehyde 5-hydroxylase (F5H; CYP84A1) are responsible for the subsequent *meta*-hydroxylations of benzene ring. C3’H catalyzes the first *meta*-hydroxylation reaction, necessary for the synthesis of both G and S-lignin units (Franke et al. 2002). On the other hand, F5H catalyzes the second *meta*-hydroxylation reaction, leading to the formation of S-lignin subunits (Supplemental Fig. S1) (Humphreys et al. 1999). Uniquely, the characterized F5H in *Selaginella moellendorffii* (SmF5H) is evolutionally distant from its angiosperm counterpart and belongs to a phylogenetically new P450 family, CYP788 (Weng et al. 2008b). Unlike angiosperm F5H, SmF5H is essentially a bifunctional *meta*-hydroxylase capable of catalyzing both 3- and 5-hydroxylations of phenylpropanoids (aldehydes and alcohols), indicating its independent evolution (Weng et al. 2010). Nevertheless, similar to its angiosperm counterpart, SmF5H can restore S-lignin biosynthesis when expressed in the *Arabidopsis* F5H-deficient mutant *fah1* (Weng et al. 2008b).

The catalysis of P450 enzymes requires redox partner(s) to supply reducing power, specifically electrons from pyridine dinucleotide cofactors such as NADPH and/or NADH (Hannemann et al. 2007). In eukaryotic cells, there are two endoplasmic reticulum (ER) electron transfer systems: the NADPH-cytochrome P450 oxidoreductase (CPR) chain and the NADH-cytochrome *b_5_* reductase (CBR)-cytochrome *b_5_* (CB5) chain. These systems support the activities of P450 enzymes and other oxidases (Liu 2022; Porter 2002; Schenkman and Jansson 2003; Kandel and Lampe 2014). In some cases, CB5 also functionally associates with CPR and accepts electrons from it to regulate P450 enzymatic activity (Bhatt et al. 2017; Zhao et al. 2023). CB5 represents an ancient heme-containing protein that is commonly found in mammals, plants, yeasts, and even purple phototrophic bacteria (Schenkman and Jansson 2003). In the yeast genome, there is only a single copy of the gene (Schenkman and Jansson 2003). while vertebrates have two isoforms, one anchored to the ER membrane and the other located at the outer mitochondrial membrane (Cowley et al. 2005; Parthasarathy et al. 2011). In contrast, flowering plant genomes often contain multiple *CB5* genes (Fukuchi-Mizutani et al. 1999; Kumar et al. 2006; Maggio et al. 2007; Smith et al. 1992; Napier et al. 1995; Hwang et al. 2004; Kumar et al. 2012; Gou et al. 2019). For instance, the model plant *Arabidopsis thaliana* genome has five annotated *CB5* genes, encoding isoforms AtCB5A (At1g26340), AtCB5B (At2g32720), AtCB5C (At2g46650), AtCB5D (At5g48810), and AtCB5E (At5g53560), respectively. These isoforms share only 40∼70% amino acid sequence identities, implicating their functional diversification.

Indeed, we previously discovered that *Arabidopsis* CB5 family member AtCB5D specializes as an indispensable electron shuttle protein, supporting the AtF5H1-catalized benzene ring 5-hydroxylation required for the synthesis of 5-hydroxylated phenolics and S-lignin monomers (Gou et al. 2019). None of the other CB5 isoforms in *Arabidopsis* can substitute for CB5D in augmenting AtF5H1 catalysis, highlighting the highly specified CB5 family in higher plants (Gou et al. 2019; Zhao et al. 2023). Moreover, the recognized CB5D member accepts electrons from both NADPH-CPR and NADH-CBR electron transfer systems and functionally associates exclusively with AtF5H1, but not C4H (and C3’H), in monolignol biosynthesis (Zhao et al. 2023). Unlike C4H and C3’H that are believed to have diverged from ancient primary metabolic cousins and evolutionarily emerged at a very early stage of land plant evolution (Werck-Reichhart and Feyereisen 2000; Alber et al. 2019; Renault et al. 2017). the invention of F5H in angiosperms has been regarded as a recent evolutionary event, coinciding with the innovation of S-lignin branch (Weng et al. 2008b; Weng et al. 2010). The identification of CB5D, tightly functionally associated with F5H in angiosperm, raises an intriguing question: did CB5D co-evolve with F5H and acquire its functional specification during the rise of S-lignin? Furthermore, similar to angiosperm F5H, SmF5H in *Selaginella* also catalyzes S-monomer biosynthesis. It remains unclear whether this independently evolved P450 enzyme also requires CB5 as redox partner for its catalysis, as its angiosperm counterpart does.

In this study, through in planta genetic exploration and heterologous whole-cell biocatalytic assays, we have discovered that CB5 family has substantially expanded and diversified across land plant lineages, with their functional specification occurring very early in the evolution of land plants. Notably, the function of CB5D in augmenting F5H-catalyzed hydroxylation for syringyl-type of phenolic synthesis emerged in the ancient embryophytes liverworts and mosses, long before the invention of angiosperm F5H and S-lignin biosynthesis. Remarkably, this early specified CB5D function has even been retained in lineages that do not produce S-lignin, such as ferns and gymnosperms. On the other hand, although S-lignin was invented in the most basal vascular plant lycophyte *Selaginella*, neither of its two CB5 orthologs has maintained CB5D function. Furthermore, unlike AtF5H1, SmF5H, similar to the monolignol biosynthetic C4H and C3’H, recruits CPR but not CB5 as its electron donor protein. These findings suggest that the P450 monooxygenase F5H in higher plant has co-opted the anciently evolved electron transfer protein CB5D for its newly invented benzene ring 5-hydroxylation.

## Results

### CB5 is expanded and diverged in land plant lineages

Among five canonical CB5 members found in *Arabidopsis*, only CB5D functions as an indispensable electron donor to support AtF5H1-catalyzed 5-hydroxylation of benzene ring for syringyl type of phenolics biosynthesis, which implicates a functional specialization of plant CB5 proteins. This functional divergence inspired us to explore deeply the evolution of plant CB5 family, particularly in respect to the emergence of the CB5D functionality in supporting F5H-catalyzed S-lignin biosynthesis.

Through blast searching using *Arabidopsis CB5D* gene as the initial query, we searched widely the available genomic and/or transcriptomic sequences of 16 representative plant species for its homologs. The selected species are across the entire green lineages, including green algae (*Volvox carteri* and *Chlamydomonas reinhardtii*), bryophytes common liverwort (*Marchantia polymorpha)* and moss (*Physcomitrella patens*), lycophyte spike moss (*Selaginella moellendorffii*), monilophytes maidenhair fern (*Adiantum aleuticum*), giant fern (*Angiopteris evecta*) and aquatic floating fern (*Salvinia_cucullata*), gymnosperms ginkgo (*Ginkgo biloba*), loblolly pine (*Pinus taeda*), norway spruce (*Picea abies*) and white spruce (*Picea glauca*), and angiosperms *Amborella* (*Amborella trichopoda*), rice (*Oryza sativa*), poplar (*Populus trichocarpa*) and petunia (*Petunia axillaris*) (Fig. 1A). The obtained putative homologs were then used as queries to conduct reciprocal blast against *Arabidopsis* genome, which gave rise to a large set of plant CB5 orthologs (Supplemental Data Set 2). In contrast to the only one *CB5* gene found in yeast and green algae, the numbers of the gene in land plant species commonly range from 2 to 7 (Supplemental Data Set 2), indicating a significant gene expansion after ancestral plants switched their aquatic habitat to the terrestrial environment. Moreover, it is obvious that the more advanced plant species contained the more copies of *CB5* homologs. For example, the flowering plants *Arabidopsis*, poplar, petunia and rice typically possess 5∼7 genes, sharply contrast to the only 1 in green algae and 2 in ancient land plant liverwort. This data implies that *CB5* has undergone dramatic expansion and diversification along land plant evolution and speciation.

**Figure 1.**
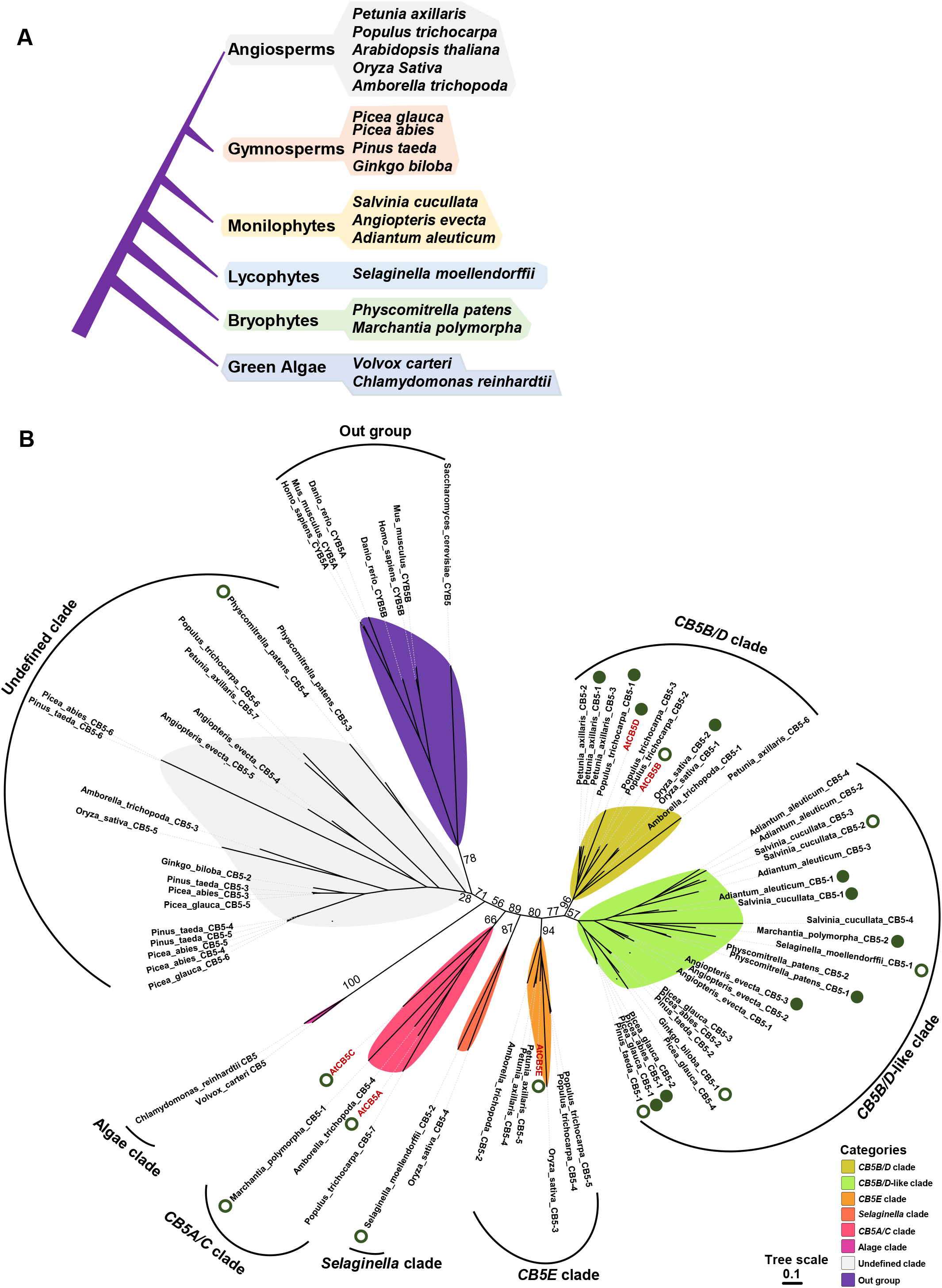
Phylogeny of CB5 homologs from representative plant taxa. **A**) A simplified version of phylogenetic tree showing the phylogenetic relationships of the plant species where the *CB5* homologs were identified. **B**) The reconstructed phylogeny of CB5 family with 71 homologs from plant species indicated in **(A**) and 7 homologs from *S. cerevisiae, H. sapiens, M. musculus* and *D. rerio*, respectively, as outgroup. Protein and CDS sequences of candidate CB5s were obtained using AtCB5D as the query via BLASTp against UCSC Genome Browser (*H. sapiens*, *M. musculus* and *D. rerio*), Saccharomyces Genome Database (*S. cerevisiae*), Phytozome (*P. trichocarpa*, *O. sativa*, *A. trichopoda*, *S. moellendorffii*, *M. polymorpha, P. patens, V. carteri* and *C. reinhardtii*), PLAZA (*P. glauca*, *P. abies*, *P. taeda* and *G. biloba*), ONEKP (*A. evecta* and *A. aleuticum*), FernBase (*S. cucullata*) and Solanaceae Genomics Network (*P. axillaris*). Maximum likelihood tree was constructed with IQ-tree2 algorithm (TIM2e+I+R4 model, 1000 SH-aLRT test and 1000 ultrafast bootstrap). Only major bifurcate bootstrap supports are marked on the main branches. *Arabidopsis CB5* genes are highlighted in magenta. The solid and open green dots indicate the genes rescued and failed in restoring *cb5d-1* defects in complementation assays, respectively. The complementation assay results of *AtCB5A*, *AtCB5B*, *AtCB5C* and *AtCB5E* (Zhao et al., 2023) were also included here.

To determine the evolutionary relationship of plant CB5 family, phylogeny reconstruction was conducted with the total of 78 *CB5* homologs found from 17 green plants and 4 outgroup species via maximum likelihood method (Fig 1B, Supplemental File 1 and 2). The constructed phylogenic tree showed a topology that all green lineage *CB5s* are distinct from their yeast and mammalian counterparts and are further clustered into several distinct clades with high bootstrap value support (Fig. 1B). Among them, the algae *CB5s* are grouped as an independent clade distinct from all other homologs of land species, suggesting substantial evolution and development of CB5s occurred along plant terrestrialization. Referenced with *Arabidopsis CB5*s classification, the homologs of land plant species are divided into highly supported clades of *CB5A/C*, *CB5B/D*, *CB5B/D*-like, *CB5E*, and a *Selaginella* clade. In addition, several diverse *CB5* homologs across entire embryophytes form a loosely positioned clade in the phylogenetic tree (undefined clade) (Fig. 1B). The *CB5A/C* clade rooted from algae counterparts likely represents the most primitive group of *CB5s* of land plants. It is composed of *CB5s* from the basal bryophyte liverwort, the basal flowering species *Amborella*, and the higher plant poplar and *Arabidopsis*. The ancestral feature of this clade of genes is reflected with the observation that *Arabidopsis* CB5A localizes in the chloroplast (Maggio et al. 2007), an organelle is thought to have evolved from endosymbiotic cyanobacteria. Consistent with previous analysis (Gou et al. 2019), *AtCB5B* and *AtCB5D* are clustered together, and with other genes exclusively from flowering lineages, constituting a unique flowering plant *CB5* clade, named as *CB5B/D* clade. This clade might represent the most advanced group of *CB5* in evolution (Fig. 1B). Being a sister to the *CB5B/D* clade is the group of *CB5* homologs from diverse land species spanning from the basal bryophyte to lycophyte, monilophyte and gymnosperm, termed as *CB5B/D*-like clade. *CB5B/D* and *CB5B/D*-like clades are positioned as a monophyletic group, suggesting their origination from a common ancestral progenitor. Notably, the homologs from the early emerging bryophytes liverwort and moss are clustered within the *CB5B/D*-like clade, hinting at the early invention of *CB5D*-like gene prior to the split and speciation of the early land plant lineages. On the other hand, the *CB5B/D* or *CB5B/D*-like sequences are retained across over the descendent tracheophytes, suggesting their high degree of conservation.

Within the phylogeny, two liverwort *CB5s* (*M. polymorpha CB5-1, MpCB5-1* and *M. polymorpha CB5-2, MpCB5-2*) are classified into evolutionarily distant clades *CB5A/C* and *CB5B/D*. Similarly, four *CB5* members of bryophyte moss (*P. patens*) are separated as two pairs and placed within two distinct clades, with one pair grouped within the *CB5B/D*-like clade and adjacent with liverwort *MpCB5-2,* and the other pair casually clustered with fern, gymnosperm and angiosperm homologs (in the undefined clade). These data indicate that *CB5* gene has undergone early gene duplication and potential functional divergence in the basal land plants.

### Complementation of *atcb5d* deficiency by *CB5* orthologs of land plant species

The emergence of *CB5B/D*-like sequences in the primitive land plant lineages implies that the functions analogous to AtCB5D might have already evolved in the ancient land species. To unequivocally demonstrate the emergence of *bona fide* CB5D in land plants, we selected 14 *CB5* homologs from *CB5B/D* and *CB5B/D*-like clades of the examined species based on their relatively high hit score with *AtCB5D* in BLASTp (Fig. 2, Supplemental Data Set 3) to conduct complementation assay to *Arabidopsis cb5d-1* mutant deficient in S-lignin and sinapoyl esters biosynthesis (Gou et al. 2019). In addition, three *CB5* homologs outside the *CB5B/D*-like clade and sharing relatively low amino acid sequence similarities with *AtCB5D*, i.e., the liverwort *MpCB5-1*, *P. patens CB5-4* (*PpCB5-4*) and *S. moellendorffii CB5-2* (*SmCB5-2*), were also included for comparation (Fig. 2, Supplemental Data Set 3).

**Figure 2.**
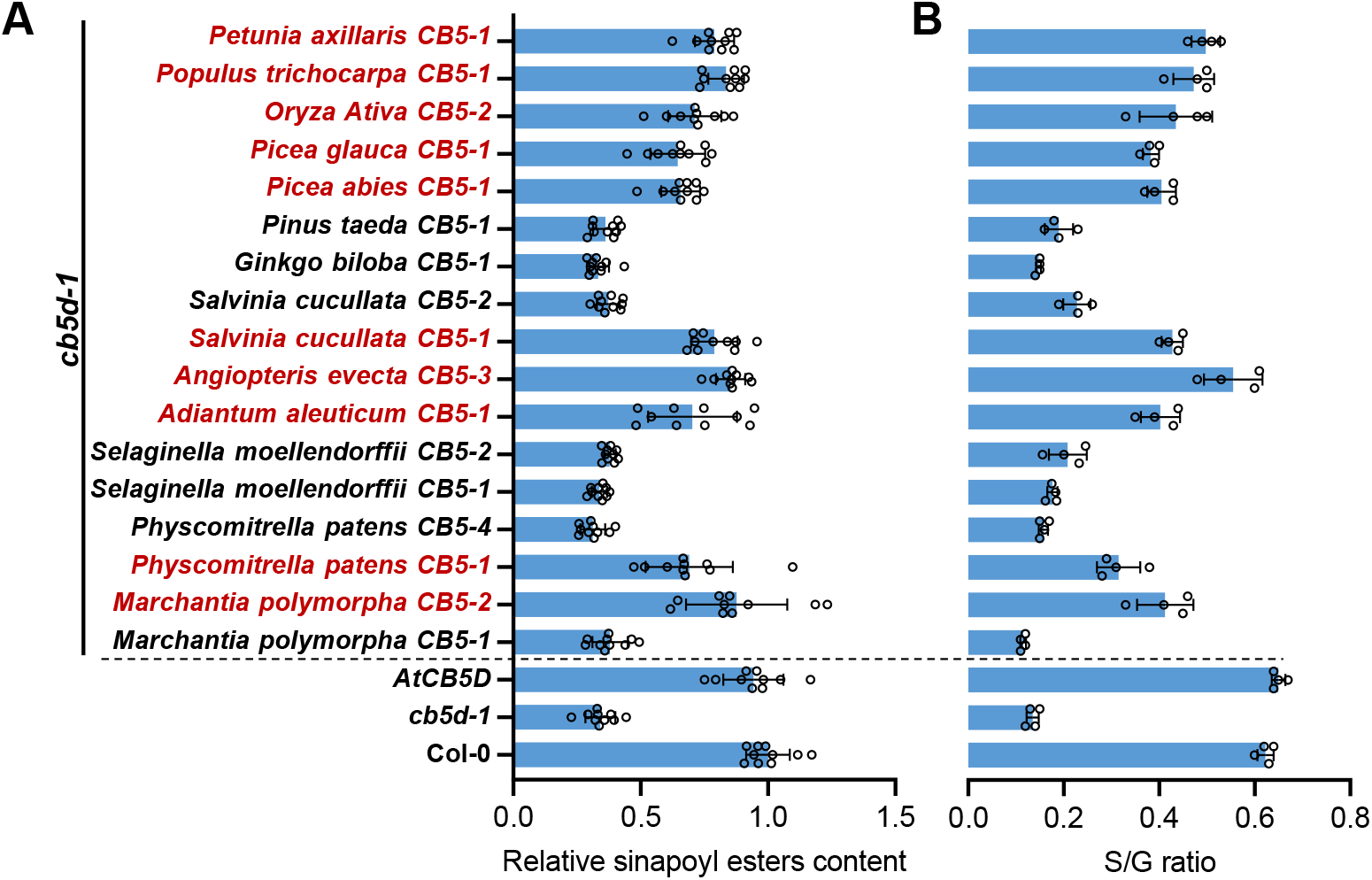
Complementation of *cb5d-1* with *CB5D* orthologs. **A**) Relative sinapoyl esters contents of the leaves and **B**) S/G ratio of stem lignin of the primary transgenic lines of the selected 17 *CB5D-like* orthologs in *Arabidopsis cb5d-1* background. One rosette leaf from 4-week-old individual primary transgenic line was sampled and every two of them were mixed representing one biology replicate for sinapoyl esters analysis. Two to four 11-week-old stems of the primary transgenic lines were pooled representing one biology replicate for lignin thioacidolytic analysis. Data are presented as means ± s.d. of ten biological replicates for sinapoyl esters quantification and four biological replicates for lignin thioacidolysis analysis. The *CB5D* orthologs that complemented *cb5d-1* are highlighted in magenta and the genes failed in complementation are labeled in black.

All 17 synthesized *CB5* genes, driven by an *Arabidopsis C4H* promoter, were respectively transformed to *cb5d-1* mutant. The *CB5* transgenes and their expression were confirmed by genome PCR in primary transgenic lines and/or by RT-qPCR in T2 seedlings (Supplemental Fig. S2). Subsequently, the leaf sinapoyl esters content and stem S-lignin (expressed as S/G ratio) were determined in the obtained transgenic lines along with the wild-type (WT) and *cb5d-1* as the controls. The quantifications were used as proxies to measure the extent of complementation of those *CB5* orthologs to the defects of *cb5d-1* in phenolic synthesis. Relative to the *cb5d-1* and WT plants, the analyzed transgenic lines were obviously displayed as two groups. Group I included *cb5d-1* mutant and the transgenic lines harboring individual *CB5* homolog *MpCB5-1, PpCB5-4, SmCB5-1* and *CB5-2, S. cucullata CB5-2, G. biloba CB5-1,* and *P. taeda CB5-1,* where their S-lignin and leaf sinapoyl ester contents remained at the similar levels as those of *cb5d-1*, indicative of the failure in complementation of *cb5d-1.* Group II encompassed WT, *AtCB5D/cb5d-1* complementation lines, and the transgenic lines harboring the rest individual *CB5* homologs of land plant species. The S/G ratio and sinapoyl ester contents in those transgenic lines were restored to the levels of the WT and *AtCB5D/cb5d-1* line, indicating successful restoration of *cb5d-1* defects with the selected *CB5* homologs from different land plant species (Fig. 2). Most interestingly, although no lignin was invented in non-vascular plants (Espineira et al. 2011), *CB5* homologs of bryophyte liverwort *MpCB5-2* and moss *PpCB5-1* effectively rescued the defects of *cb5d-1*, strongly evident that the *bona fide* CB5D has been invented in the ancient embryophytes prior to the occurrence of vascular plants (Fig. 2). By contrast, the paralogs *MpCB5-1* of liverwort and *PpCB5-4* of moss failed in restoring the *cb5d-1* defects (Fig. 2), suggesting the functional divergence and specialization of CB5 family in the basal land plants.

Furthermore, the CB5D function was retained in descendent euphyllophytes (including ferns, gymnosperms, and angiosperms). The homologs from ferns (*S. ucullate CB5-1, A. evecta CB5-3 and A. aleuticum CB5-1),* pines (*P. glauca CB5-1* and *P. abies CB5-1*), and angiosperms rice (*O. sativa CB5-2*), poplar (*P. trichocarpa CB5-1*) and petunia (*P. axillaris CB5-1*) all restored *cb5d-1* defects in sinapoyl esters and S-lignin biosynthesis (Fig. 2). These data indicate that the early emerged CB5D function was inherited and sustained in the higher plants during evolution.

### Liverwort MpCB5-2 effectively supports *Arabidopsis* F5H catalysis

The restoration of *cb5d-1* with liverwort *MpCB5-2* transgene encouraged us to further dissect the biochemical properties of liverwort *CB5* homologs. When fused with yellow fluorescent protein (YFP) and transiently expressed in tobacco leaf epidermal cells, two liverwort CB5s showed distinct subcellular localization patterns. Consistent with phylogenetic classification in *CB5A/C* clade, the fluorescence distribution of YFP-MpCB5-1 fusion appeared in the chloroplast (Fig. 3A), which is similar to that reported on AtCB5A; whereas MpCB5-2 that rescues *cb5d-1* defects displayed a typical ER network localization pattern, implicating its potential function as an ER-electron transfer chain component (Fig. 3B). The recombinant MpCB5 proteins with truncation of their transmembrane domains were then produced and purified from *E.coli*. Both recombinant MpCB5-1 and MpCB5-2 could be reduced by reducing agent dithionite and displayed typical cytochrome *b_5_* spectroscopic characteristics with a maximal soret absorption at 412 nm in their oxidized form and at 423 nm along with addition absorption at 528 and 557 nm when they were completely reduced (Fig. 3C and 3D) (Zhao et al. 2023). When *Arabidopsis* CBR1 and CPR (namely ATR2) were used to reduce the recombinant MpCB5s, MpCB5-2, the CB5B/D-like protein, was more effectively reduced compared to MpCB5-1 (Fig. 3E and 3F), implicating its better intrinsic electron accepting ability.

**Figure 3.**
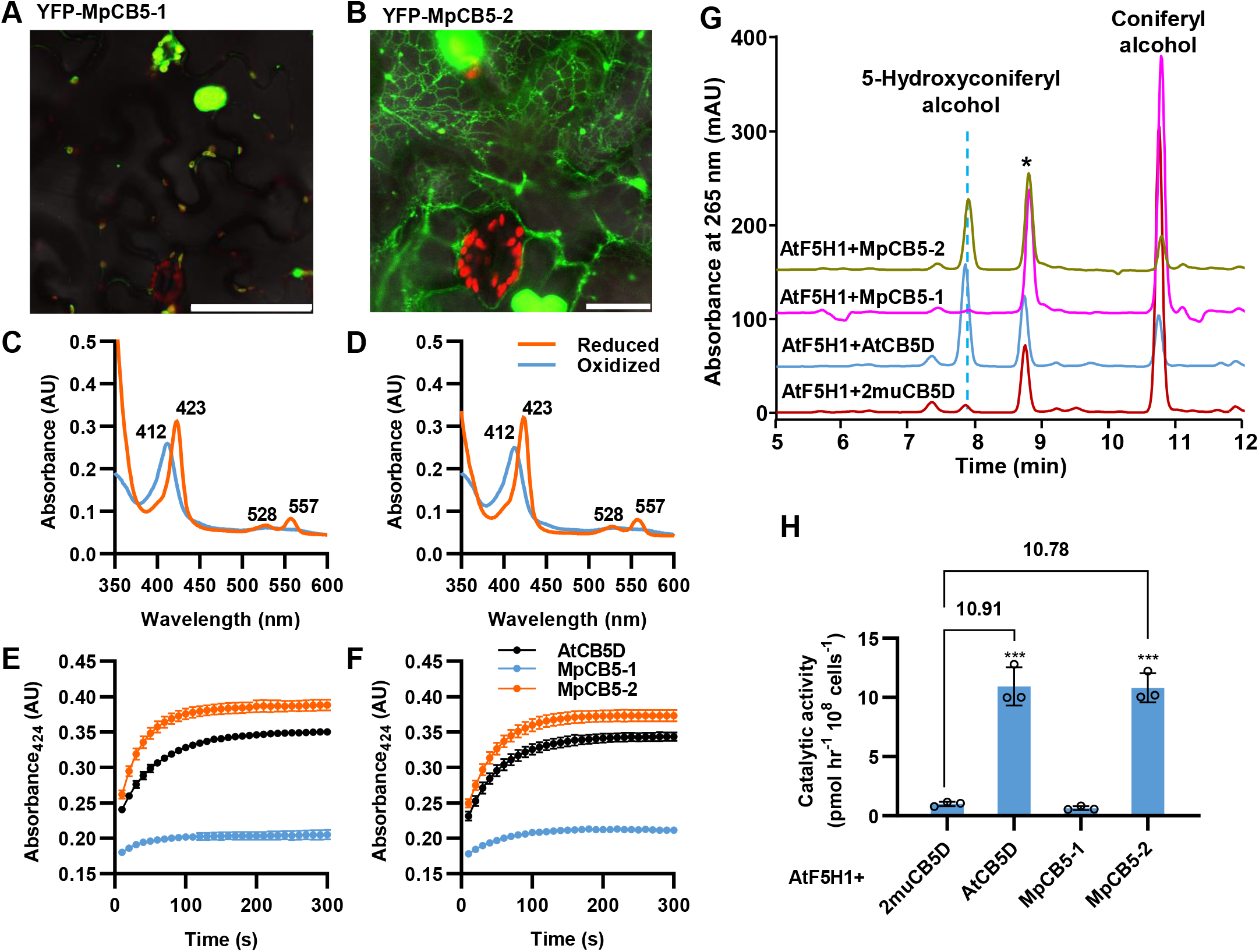
Characterization of liverwort *MpCB5s.* **A** and **B)** Fluorescence distribution of YFP-MpCB5-1 fusion **(A)** and YFP-MpCB5-2 fusion **(B)** in *N. benthamiana* leaf epidermal cells. Green pseudo color indicates YFP fluorescence signals, and red color indicates chloroplast autofluorescence. Scale bars, 100 µm **(A)** and 20 µm **(B)**. **C** and **D)** UV-visible absorbance spectra of the recombinant MpCB5-1 **(C)** and MpCB5-2 **(D)** in the oxidized (blue) and reduced (orange) status. The recombinant proteins (7µM) were reduced by dithionate at room temperature. **E** and **F)** Changes of absorbance at 424 nm of MpCB5s over the indicated period of reaction with 7.2 nM CBR1 **(E)** or 720 nM ATR2 **(F)**. The absorption was recorded for 30 cycles with a 10 s interval. Data are presented as means ± S.D. of three independent experiments. **G)** HPLC-UV profiles of F5H-catalyzed biotransformation of coniferyl alcohol to 5-hydroxyconiferyl alcohol within *E. Coli*. The asterisk indicates a non-specific peak. **H)** The catalytic activity of different F5H-CB5 fusion proteins. Coniferyl alcohol was fed as substrate and 5-hydroxyconiferyl alcohol product was monitored by HPLC. The fold changes were shown on the solid lines, compared with the control cells devoid of functional CB5 patterner. Date are presented as means ± S.D. of three biological replicates. Asteroids indicate significant difference with \*\*\**P* < 0.001 (two-tailed Student’s t tests).

To further assess the potential functional association of liverwort CB5s with F5H, we developed a whole-cell biocatalytic assay within *E.coli* strain via co-expression of *CB5* and *F5H*. The truncated MpCB5s, AtCB5D or a mutant variant of CB5D, i.e., 2muCB5D, with removal of the transmembrane domains was co-expressed with AtF5H1 in *E.coli*. 2muCB5D bears the substitutions of two histidine residues that are crucial for its heme binding to alanine, therefore, devoid of electron transfer property (Gou et al. 2019). It was included in the assay as a control. The co-expressed two proteins were fused together via a lambda linker (GSTSSGSG) (Schuckel et al. 2012) and placed within the same expression cassette to ensure their same level of expression. After induction of protein expression with IPTG, the accumulated recombinant proteins were monitored via SDS-PAGE analysis, which showed strong Coomassie blue stained bands (Supplemental Fig. S3). Then the induced *E. coli* cells were incubated with coniferyl alcohol, the substrate of F5H, and the transformed products were monitored with HPLC and UHPLC-MS (Supplemental Fig. S4).

The AtF5H1-2muCB5D and AtF5H1-AtCB5D fusions were pre-tested to verify the reliability of the whole-cell catalytic system. Compared with 2muCB5D, the co-expression of AtF5H1 with AtCB5D resulted in up to 11-fold increase of F5H catalytic activity in converting coniferyl alcohol to 5-hydroxyconiferyl alcohol product (Fig. 3G and 3H), demonstrating that the truncated AtF5H1 and AtCB5D work properly in *E. coli* cells and also confirming that AtCB5D is indeed an effective electron donor for AtF5H1-catalyzed benzene ring 5-hydroxylation. In addition, this result also implies that the endogenous *E. coli* redox systems can effectively reduce CB5 protein, which is consistent with the previous observations (Hatakeyama et al. 2016; Ichinose et al. 2004). Next, AtF5H1-MpCB5-1 and AtF5H1-MpCB5-2 fusions were examined. Consistent with the *in planta* complementation assay (Fig. 2), MpCB5-2 substantially enhanced AtF5H1 activity to the level displayed by AtF5H1-AtCB5D combination. However, MpCB5-1 essentially showed no effect on AtF5H1 catalytic activity (Fig. 3G and 3H). These data affirm that liverwort possesses the *bona fide* CB5D protein. Furthermore, we tested whether the primordial CB5s originated from algae possess CB5D function. With synthesized *C. reinhardtii CB5* (*CrCB5*) and *V. carteri CB5* (*VcCB5*), we expressed their fusion protein with *AtF5H1* in *E. coli* and conducted the whole-cell catalytic assay. Neither of the algal CB5 proteins exhibited the functional effect on AtF5H1 catalytic activity (Supplemental Fig. S5). These assay results assert that CB5 protein functionally diverged after land plant colonization and the ancient embryophyte liverwort possesses the *bona fide* CB5D comparable to the higher plant counterpart.

### F5H is not invented in bryophyte, fern and gymnosperm

The observation of *bona fide* CB5D in the species of bryophyte, fern and gymnosperm inspired us to re-examine whether S-lignin and *F5H* like genes ever evolved in those land lineages, even though the previous studies have defined that S-lignin biosynthesis and F5H were restricted only in angiosperms and the primitive vascular plant *Selaginellales* order (Weng et al. 2008a; Weng et al. 2010). Applying conventional diagnostic thioacidolysis method that cleaves the β-aryl ester bonds of lignin polymer, we examined lignin composition in fern *A. aleuticum* and gymnosperm norway spruce (*P. abies*), both have *CB5D*-*like* genes that are able to restore *cb5d-1* defects, and loblolly pine (*P. taeda*) whose *CB5D-like* gene failed in rescuing *cb5d-1* deficiency. No detectable level of S-lignin monomers was found in all the examined plants (Supplemental Fig. S6A), inferring that the retention of CB5D in fern (*A. aleuticum*) and norway spruce (*P. abies)* is not related to the S-lignin biosynthesis. Furthermore, we re-searched the potential *F5H* homologous sequences using *AtF5H1* as the query against the genomic or transcriptomic sequences of liverwort (*M. polymorpha),* moss (*P. patens)*, and white spruce *(P. glauca)* whose genomes contain *CB5D-like* genes that rescued *cb5d-1* mutant, and *Ginkgo biloba* whose *CB5D-like* gene failed in restoring *cb5d-1* deficiency but was reported with production of S-lignin subunits in the old biochemical study (Uzal et al. 2009). Although the obtained hits showed low sequence similarity, around 34% to 44% to AtF5H1 at amino acid level, four candidate genes that are relatively more similar to *AtF5H1*, i.e., *Mapoly0025s0014.1*, *Pp3c1_1880*, *PGL00005126*, and *GBI00006168* were synthesized and transformed, under the control of *Arabidopsis C4H* promoter, to the *fah1-2* mutant that is defective in S-lignin biosynthesis due to the mutation of *AtF5H1* gene (Supplemental Fig. S6B and S6C) (Chapple et al. 1992). No 5-hydroxylated phenolics sinapoyl esters in the leaves and S-lignin monomers from the stems were recovered in their primary transformants, indicating that the selected candidates do not functionally complement *fah1* deficiency (Supplemental Fig. S6D and S6E). Therefore, consistent with the previous study (Weng et al. 2008b), it is most likely no functional *F5H* homologs ever emerged in bryophytes or gymnosperm, regardless of whether the plants possess the functional *CB5D*.

### *Selaginella* doesn’t possess the evolved functional CB5D

*Selaginella* synthesizes S-lignin in its cortex (Weng et al. 2010). An independently evolved cytochrome P450 CYP788A1, namely SmF5H, was characterized responsible for its S-lignin biosynthesis (Weng et al. 2008b; Weng et al. 2010). We aimed to explore whether this early evolved F5H of the primitive vascular plant also requires CB5 as the electron donor for its catalysis as does its counterpart AtF5H1. Searching *Selaginella* genome, two *CB5* homologs, *Sm271829* (*S. moellendorffii* CB5-1, *SmCB5-1*) and *Sm148632* (*S. moellendorffii CB5-2*, *SmCB5-2*) were revealed. In the *CB5* family phylogeny, similar to the liverwort *MpCB5s*, two *SmCB5s* were separated into two distinct clades, with *SmCB5-1* clustered with liverwort *MpCB5-2* and moss *PpCB5-1* and *CB5-2*, and grouped within *CB5B/D*-like clade, while *SmCB5-2* separated as an individual clade (Fig. 1B), which indicate their sequence and perhaps functional divergence. The encoded proteins of both *SmCB5* homologs, when fused with YFP and expressed in *N. benthamiana* leaf epidermal cells, displayed a typical ER membrane-localization pattern (Fig. 4A and B), reminiscent of AtCB5D localization (Gou et al. 2019). To verify if SmCB5s like AtCB5D that functions as electron donor for F5H-catalyzed reaction *in planta*, two *SmCB5* genes were transformed to *cb5d-1* mutant along with a set of *CB5* homologs from other species (Fig. 2) to examine their abilities in restoring the defects of sinapoyl esters and S-lignin biosynthesis of the mutant. Although both transgenes were confirmed the expression in the generated transgenic lines (Supplemental Fig. S7), under ultraviolet (UV) illumination, all the primary transgenic plants, however, as did the *cb5d-1* mutant, appeared red color in their rosette leaves, an indicator of lack or the low accumulation levels of autofluorescent phenolics (Fig. 4C) (Ruegger and Chapple 2001). Measuring the contents of leaf and seed sinapoyl esters and stem S-lignin of transgenic plants, we found that all the 5-hydroxylation-derived phenolics remained at the similar levels as those detected in *cb5d-1* mutant, confirming no restoration of syringyl type of phenolics biosynthesis in the transgenic plants harboring either *SmCB5-1* or *SmCB5-2* (Fig. 4D to 4F). These results suggest that neither SmCB5s possesses *bona fide* CB5D function that can support AtF5H1 catalysis.

**Figure 4.**
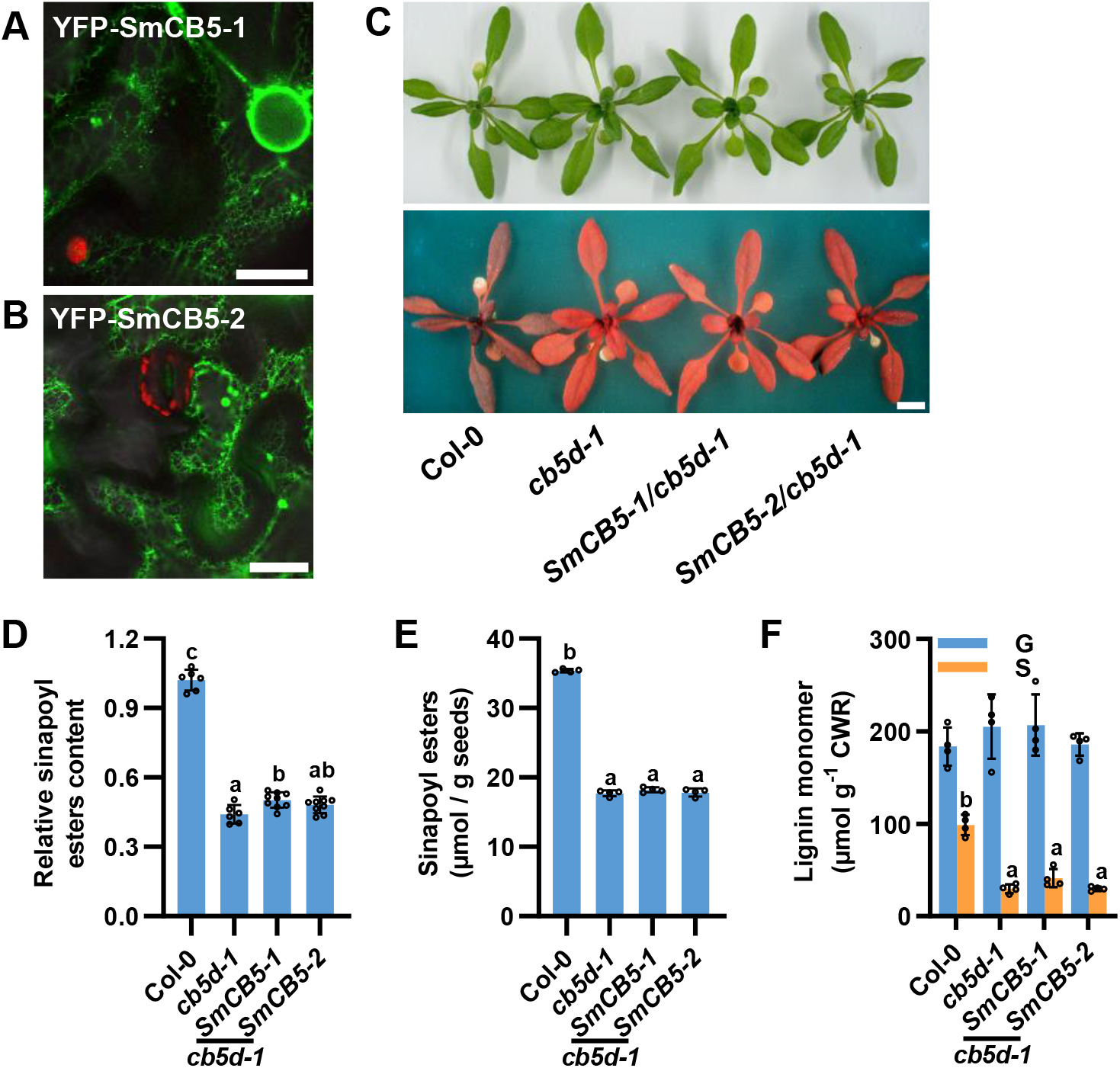
Subcellular localization and complementation assay of SmCB5s. **A** and **B)** Fluorescence distribution patterns of YFP-SmCB5-1 **(A)** and YFP-SmCB5-2 **(B)** in *N. benthamiana* leaf cells. Green color indicates YFP fluorescence signals, and red color indicates chloroplast autofluorescence. Scale bar, 20 µm. **C)** Rosette leaves of Col-0 WT, *cb5d-1* and *Arabidopsis* T2 transgenic lines of *SmCB5s* under visible light (upper) and ultraviolet light (lower). Scale bar, 1cm. **D** to **F)** Relative content of sinapoyl esters in the 1-week-old T2 seedling **D)**, sinapoyl esters content in T3 mature seeds **E)**, and lignin monomer contents released by thioacidolysis in the 11-week-old stems of primary transformants **F)**. Seedlings from one petri dish were mixed representing one biology replicate for sinapoyl esters analysis. Seeds from a single plant were pooled as one biology replicate for seed sinapoyl esters analysis. Stems from two to four lines were pooled as one biology replicate for lignin thioacidolysis analysis. CWR, cell wall residuals. Data are presented as means ± S.D. of six **(D)**, or four **(E** and **F)** biological replicates. Letters above the bars indicate significant differences (*P* < 0.05, one-way ANOVA test).

Since SmF5H evolved independently as to counterpart AtF5H1, it might be possible that SmCB5s co-evolved with SmF5H and favor to functionally associate with it instead with AtF5H1. To test this possibility, we first expressed *SmF5H* under the control of AtC4H promoter in *fah1-2* and confirmed that the expression substantially restored the accumulation of both the leaf sinapoyl esters and the stem S-lignin biosynthesis (Supplemental Fig. S8). This confirms that SmF5H indeed could function as a ferulate 5-hydroxylase in *Arabidopsis* as previously reported (Weng et al. 2008b). We then generated a homozygous *fah1-2 cb5d-2* double mutant via genetic cross (Supplemental Fig. S9) and transferred *SmF5H* alone or together with *SmCB5-1* or *SmCB5-2* into it (Supplemental Fig. S10). The *fah1-2 cb5d-2* double mutant like *fah1-2* nearly completely depleted the accumulation of syringyl type of phenolics (Fig. 5). Expression of *SmF5H* in *fah1-2 cb5d-2* partially rescued the synthesis and accumulation of leaf and seeds sinapoyl esters and stem S-lignin (Fig. 5A-D). Nevertheless, co-expression of *SmF5H* with either of *SmCB5s* in *fah1-2 cb5d-2* didn’t further enhance sinapoyl esters content and S-lignin accumulation, compared with the double mutant lines harboring the expressed *SmF5H* alone (Fig. 5). These data are evident that SmCB5 homologs are not the obligatory electron donor for SmF5H-catalyzed reaction *in planta*. Notably, compared with *AtF5H1*/*fah1-2 cb5d-2* transgenic lines, expression of *SmF5H* in the same background resulted in a discernibly higher level of restoration of the leaf and seed sinapoyl esters and stem S-lignin (Fig. 5), further suggesting that SmF5H is functionally more independent from CB5 electron donor *in planta*. Taken together, these results indicate that no functional CB5D was retained in *Selaginella* and SmF5H does not necessarily require CB5 electron donor protein for its catalysis.

**Figure 5.**
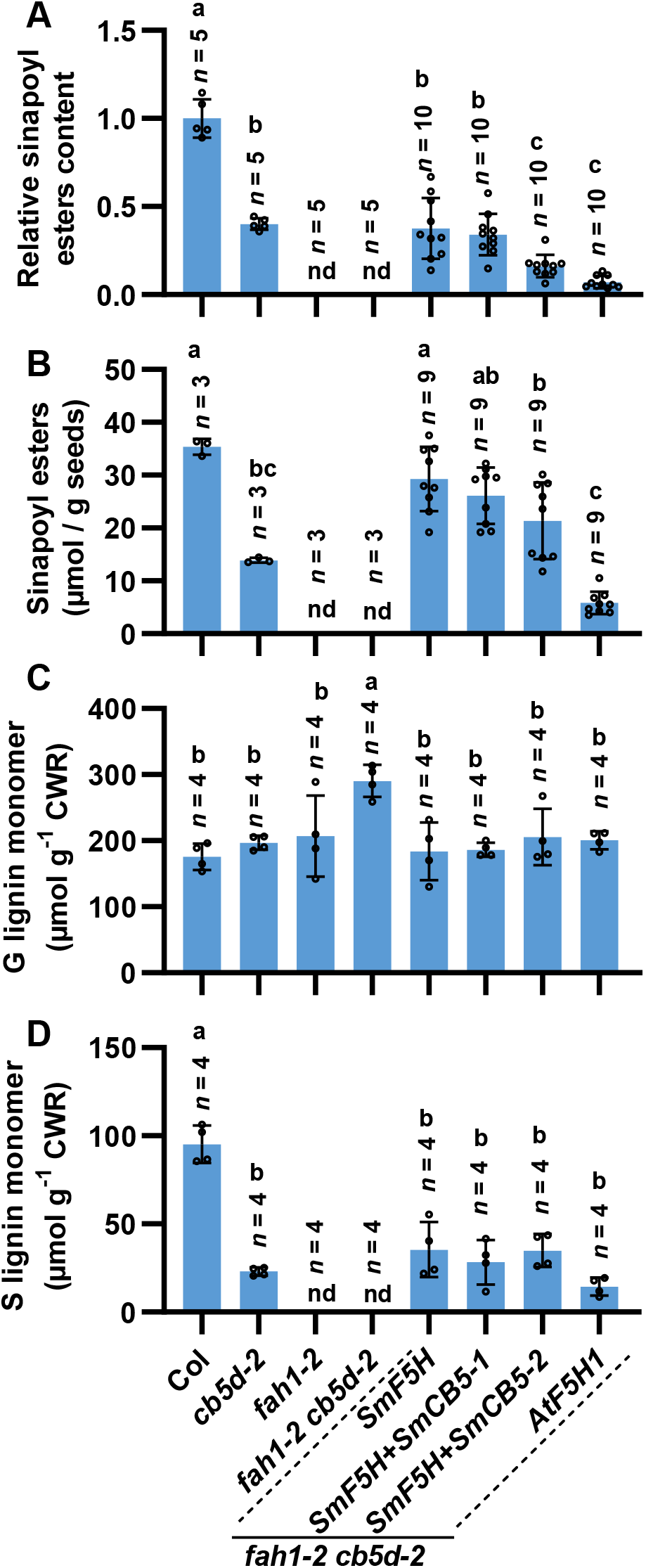
Complementation of *fah1-2 cb5d-2* mutant with *SmF5H* and *SmCB5s*. **A)** Relative contents of sinapoyl esters in the 4-week-old rosette leaves of primary transformants. One rosette leaf from a 4-week-old individual primary transgenic line was sampled and every two of them were mixed representing one biology replicate for sinapoyl esters analysis. **B)** Sinapoyl esters contents in T2 mature seeds. Seeds from a single transgenic plant were representing one biology replicate. **C** and **D)** Lignin monomer contents released by thioacidolysis analysis in the 11-week-old stems of primary transformants. Stems from two to four lines were pooled representing one biology replicate. Data are presented as means ± SD of the indicated biological replicates. Letters above the bars indicate significant differences (*P* < 0.05, one-way ANOVA test). nd, not detected. CWR, cell wall residuals.

### SmCB5s are the functional electron carriers but not required by SmF5H

The failure of *SmCB5s* in restoring *cb5d-1* deficiency in syringyl type of phenolic biosynthesis promoted us to further examine if both SmCB5s are capable of electron transfer. The recombinant SmCB5 proteins with removal of their C-terminal transmembrane domains were generated and purified from *E. coli*. Both the recombinant SmCB5s displayed a typical CB5 absorption soret band at 413 nm, indicating that they function properly. When treated with reducing agent dithionite, they were readily reduced with characteristic absorbance at 423 nm (Fig. 6A and 6B), resembling those of known CB5s (Zhao et al. 2023). When incubated with NADH and the recombinant *Arabidopsis* CBR1, both SmCB5s displayed a steady transition to the absorbance at 424 nm, indicative of an effective reduction (Fig. 6C). These data suggest that both SmCB5s can similarly accept electrons from reductant NADH. However, when NADPH and the recombinant ATR2, a CPR from *Arabidopsis,* were applied, two SmCB5 proteins displayed different reducing behaviors. SmCB5-2 like the typical CB5 proteins showed a steady switch from its oxidation status to the reduction status and eventually reached to and sustained at the maximal reduction status (Fig. 6D and 6F); whereas SmCB5-1, the CB5B/D-like protein, appeared to be rapidly reduced by ATR2 at the initial reaction stage then steadily return to its oxidized status (Fig. 6D and 6E).

**Figure 6.**
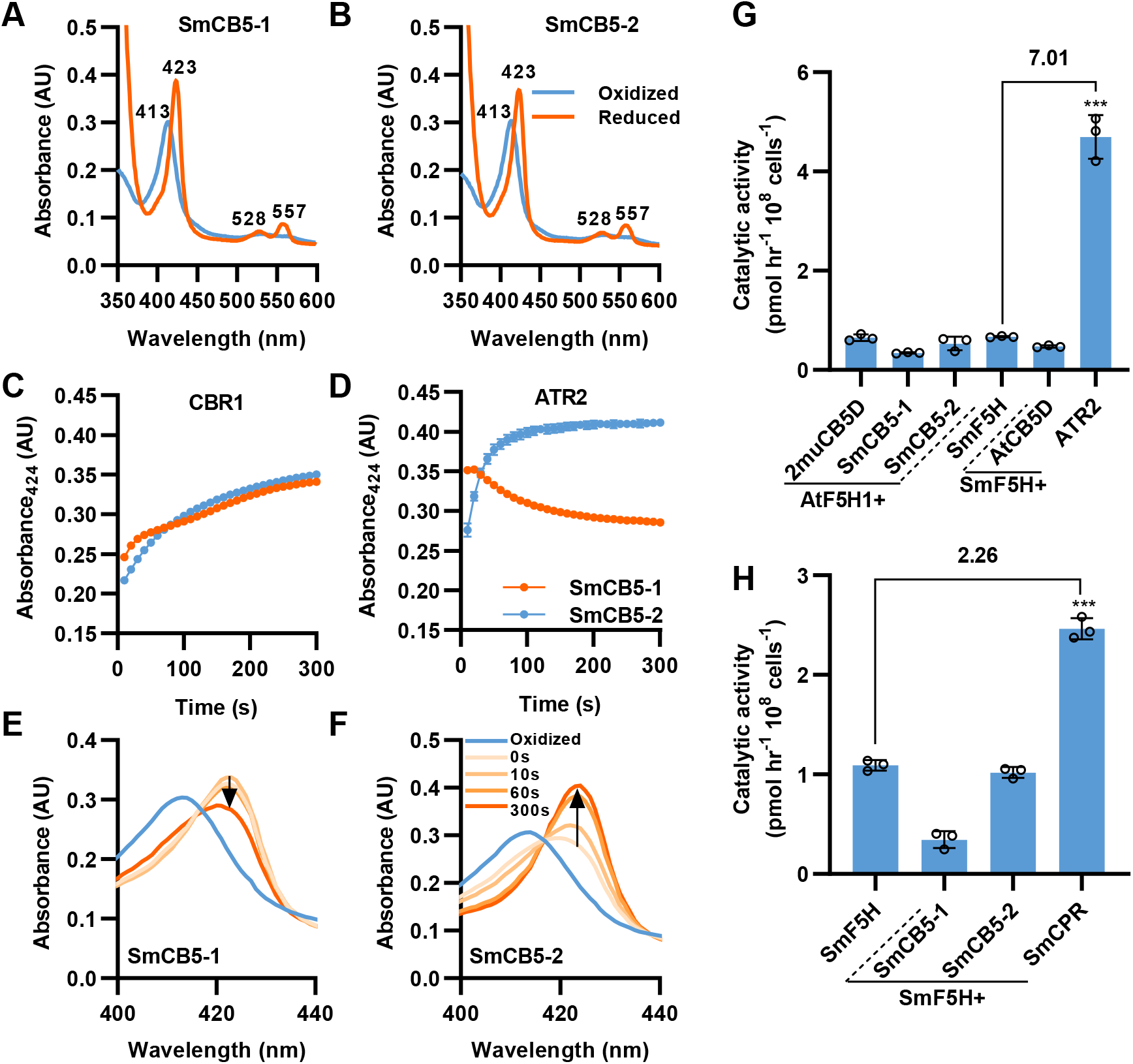
Characterization of functional properties of SmCB5s. **A** and **B)** UV-visible absorbance spectra of the recombinant SmCB5-1 **A)** and SmCB5-2 **B)** in the oxidized (blue) and reduced (orange) status. The recombinant proteins (7µM) were reduced by dithionate at room temperature. **C** and **D)** Real time changes of absorbance of SmCB5s at 424 nm over the indicated period of reaction with 7.2 nM CBR1 **(C)** and 720 nM ATR2 **(D)**. The absorption was recorded for 30 cycles with a 10 s interval. Data are presented as means ± SD of three independent experiments. **E** and **F)** UV-visible absorbance spectra of SmCB5-1 **(E)** and SmCB5-2 **(F)** in the oxidized (prior to the addition of NADPH and ATR2) and reduced (after the addition of NADPH and ATR2 for 0, 10, 60 and 300 s) status. Three repeating assays were performed with the same results; and one was presented. Note that the arrow points out the direction of the change of maximum sorbet absorption. **G** and **H)** The catalytic activities of different F5H-CB5 fusion proteins in *E. coli* whole cell assays. Coniferyl alcohol was fed as substrate and 5-hydroxyconiferyl alcohol product was monitored by HPLC. The fold changes are shown on the solid lines compared with the control cells devoid of CB5 patterner. Date are presented as means ± SD of three biological replicates. Asterisks indicate significant difference with \*\*\**P* < 0.001 (two-tailed Student’s *t* tests).

To obtain a better understanding of the electron transfer capacity of SmCB5s to F5H, we applied *E. coli* whole-cell biocatalytic system. SmCB5-1 or SmCB5-2 was first fused with AtF5H1 and co-expressed; neither co-expression enhanced AtF5H1-catalytic activity (Fig. 6G), which strongly contrasted to the substantial enhancement of AtF5H1 activity via co-expression with AtCB5D in the same assay system (Fig. 3H). On the other hand, SmF5H was also fused with *Arabidopsis* electron transfer component AtCB5D or ATR2. The co-expression with AtCB5D did not alter the performance of SmF5H. By contrast, when ATR2 was co-expressed, the catalytic activity of SmF5H was enhanced up to 7-fold (Fig. 6G), strongly suggesting that SmF5H, like *Arabidopsis* C4H and C3’H (Gou et al. 2019; Zhao et al. 2023), relies on CPR redox partner only. Next, SmF5H was co-expressed with SmCB5-1, SmCB5-2 or SmCPR, respectively. Again, neither SmCB5s improved the catalytic efficiency of SmF5H, but SmCPR showed substantial enhancement on the catalytic activity of SmF5H (Fig. 6H). All these results are strongly evident that SmF5H relies on CPR-mediated electron transfer system, and CB5 is unnecessary for SmF5H catalysis, although both SmCB5 proteins are the functional electron carriers.

### The helix 5 of CB5 heme binding domain influences its functional specialization

Although neither SmCB5 likes AtCB5D and supports F5H catalysis, it is notable that *SmCB5-1* phylogenetically clusters with *MpCB5-2* and *PpCB5-1* and grouped within *CB5B/D*-like clade (Fig. 1B); and both MpCB5-2 and PpCB5-1are the proven *bona fide* CB5Ds (Fig. 2). This phylogenetic relationship appears to suggest that SmCB5-1 should have the evolved *bona fide* CB5D functionality as did its evolutionary sister MpCB5-2. This paradox motivated us to explore the underlying reasons that caused the loss of CB5D function of SmCB5-1. Homolog modeling-based structural determination via AlphaFold-2 (Jumper et al. 2021) on AtCB5D and SmCB5-1 revealed a similar overall structure with five α-helices constituting their cytochrome *b_5_* domains; and the helices 2 to 5 (H2 to H5) encompass the heme-binding domain (Supplemental Fig. S11A and S11B). Comparing CB5 orthologs that show *bona fide* CB5D function in the *cb5d-1* complementation assay (classified as Group 1) with those showing no CB5D function (Group 2) (Supplemental Fig. S11C), we found that within the cytosolic heme-binding domain, the two functionally distinct groups of CB5 proteins differ mostly in their amino acid residues of H5 (Fig. 7A and Supplemental Fig. S11C). Among a total of 10 amino acid residues within the H5 fragment, we observed that the prominent differences between two groups of CB5s were the substitutions of negatively charged acidic amino acid residues with the uncharged or positively charged residues (Fig. 7A and Supplemental Fig. S11C). H5 in Group I contains the conserved X7-D/E-E/D-X sequence, designated as H5-D/E motif, where X represents non-negatively charged amino acid residues (Fig. 7A); whereas the acidic residues (Asp and/or Glu) were frequently substituted in Group 2 of CB5 proteins.

**Figure 7.**
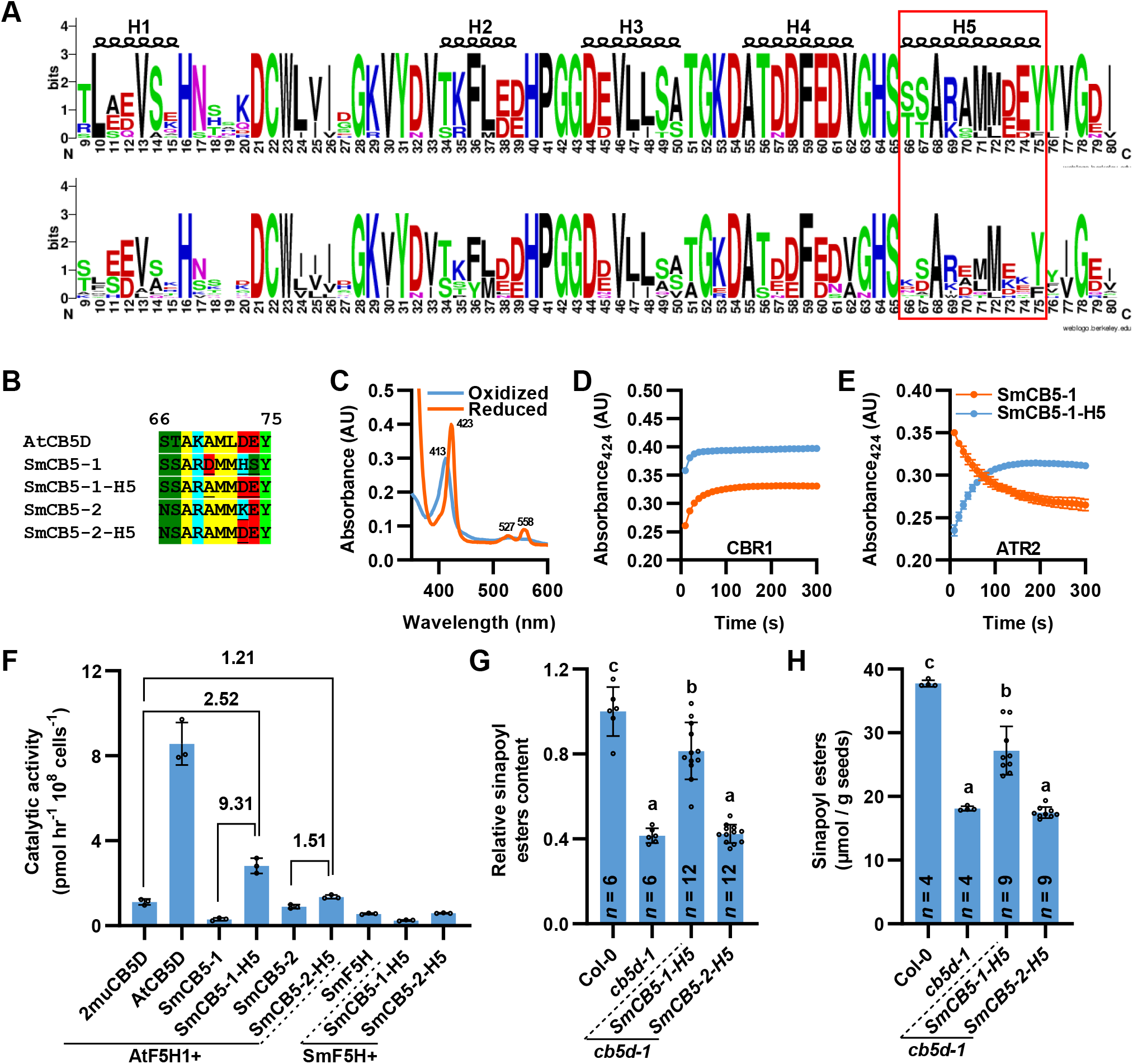
Characterization of the functional influence of helix 5 of SmCB5-1. **A)** WebLogo presentation of cytochrome *b_5_* domain of Group I (upper) and Group II (lower) CB5s. The helix 5 (H5) portion was boxed with a red rectangle. **B)** The amino acid sequences of the H5 region of AtCB5D, SmCB5s, and the created SmCB5 mutant variants. **C)** UV-visible absorbance spectra of the mutant variant SmCB5-1-H5 in its oxidized (blue) and reduced (orange) status. The recombinant CB5 protein (7µM) was reduced by dithionate at room temperature. **D** and **E)** Reduction of the recombinant SmCB5-1 and SmCB5-1-H5 by CBR1 **(D)** and ATR2 **(E)**. 7 µM recombinant protein was incubated with 7.2 nM CBR1 **(D)** or 3.6 µM ATR2 **(E)** at room temperature and the reduction reaction was initiated by the addition of either 100 µM NADH **(D)** or 100 µM NADPH **(E)** and the absorbance at 424 nm were recorded with a microplate reader for 30 cycles with 10 s interval. **F)** The catalytic activity of different F5H-CB5 fusion proteins in *E. coli* whole cell biocatalytic assay. Coniferyl alcohol was fed as substrate and 5-hydroxyconiferyl alcohol product was monitored by HPLC. The fold changes are shown on the solid lines compared with the indicated control cells. **G** and **H)** Relative content of sinapoyl esters in the 4-week-old leaves of primary transgenic lines harboring *SmCB5-1-H5* or *SmCB5-2-H5* **(G)** and sinapoyl esters content in their T2 mature seeds **(H)**. Rosette leaves from two individual transgenic lines were pooled representing one biology replicate for leaf sinapoyl ester analysis. Seeds from individual line were collected representing one biology replicate for seed sinapoyl esters analysis. Data are presented as means ± SD of indicated biological replicates. Letters above the bars indicate significant differences (*P* < 0.05, one-way ANOVA test).

Specifically, the negatively charged Asp73 and Glu74 in AtCB5D were respectively replaced with His and Ser in SmCB5-1; conversely, Asp70 in the H5 of SmCB5-1 was replaced with the uncharged Ala in AtCB5D (Fig. 7B). In addition, the negatively charged Asp73 of AtCB5D was replaced with positively charged Lys in the corresponding position of SmCB5-2 (Fig. 7B). To assess whether these substitutions affect the functionality of SmCB5s as a *bona fide* CB5D, we generated mutant variants from CB5B/D-like SmCB5-1 with triple mutations D67A/H70D/S71E, named SmCB5-1-H5, and from non-CB5B/D-like SmCB5-2 with K81D, designated SmCB5-2-H5. Prior to the *in planta* test, the truncated SmCB5-1-H5 variant with removal of its transmembrane domain was expressed and purified from *E. coli* and validated spectroscopically for its CB5 protein properties. The oxidized and dithionite-reduced SmCB5-1-H5 exhibited typical CB5 characteristics (Fig. 7C). When SmCB5-1-H5 was incubated with CBR1 and NADH, it was more efficiently reduced compared to its parental SmCB5-1, implicating a higher electron transfer capacity (Fig. 7D). Moreover, upon incubation with ATR2 and NADPH, SmCB5-1-H5 began to display a typical CB5 reducing pattern as that of AtCB5D, which sharply contrasted to its parental SmCB5-1 protein (Fig. 7E). Subsequently, we co-expressed the SmCB5 mutant variants with AtF5H1 or with SmF5H in *E. coli*, respectively. Compared to the parental SmCB5-1, co-expression of SmCB5-1-H5 with AtF5H1 increased the P450 catalytic activity more than 9-fold, suggesting that the mutations with acidic amino acid residues in H5 confer *bona fide* CB5D activity. Additionally, co-expression of SmCB5-2-H5 with AtF5H1 also slightly elevated P450 activity by 1.5-fold compared to its parental protein, the non-CB5B/D-like SmCB5-2 (Fig. 7F). Nevertheless, neither SmCB5 variants enhanced SmF5H catalytic activity when they were co-expressed in *E. coli.* These results further confirm that SmF5H does not necessarily require CB5 as redox partner for its catalysis. Next, we transformed the genes encoding both mutant variants into *Arabidopsis cb5d-1* (Supplemental Fig. S12). Measuring sinapoyl esters contents in the leaves and seeds of transgenic lines, we found that *SmCB5-1-H5* but not *SmCB5-2-H5* substantially complemented the *cb5d-1* deficiency (Fig. 7G and 7H). These data suggest that evolving and preserving H5-D/E motif in the CB5D-like protein is vital to confer its electron donor function in supporting F5H catalysis. On the other hand, for non-CB5B/D-like proteins, apart from H5-D/E motif, additional sequence element(s) contributes to their functional specialization.

By searching and comparing the occurrence of helix 5 sequences in green lineages that have available genomic or transcriptomic sequences, we counted the numbers of plant species that possess at least one of the recognized H5-D/E motif. Notably, all the examined dicots and 27 out of 28 monocots retained the sequence containing H5-D/E motif (Fig. 8A). The one examined monocot species, *Phalaenopsis equestris*, that missed the motif might be due to its low quality of genomic sequencing. Among bryophytes, monilophytes, and gymnosperms, not all the species possessed the sequence of H5-D/E motif, implicating that the early evolved *CB5D* gene might be lost in some of the species during evolution. Interestingly, while none of *Selaginellale* species held H5-D/E motif, all the species in other two orders of lycophytes, *Isoetales* and *Lycopodiales*, retained the sequence encoding this motif. Finally, none of algal species was found to have the characteristic H5-D/E motif, consistent with their functional validation (Supplemental Fig. S5). These findings underscore the significance of helix 5 in conferring and sustaining the functional evolution of a *bona fide* CB5D.

**Figure 8.**
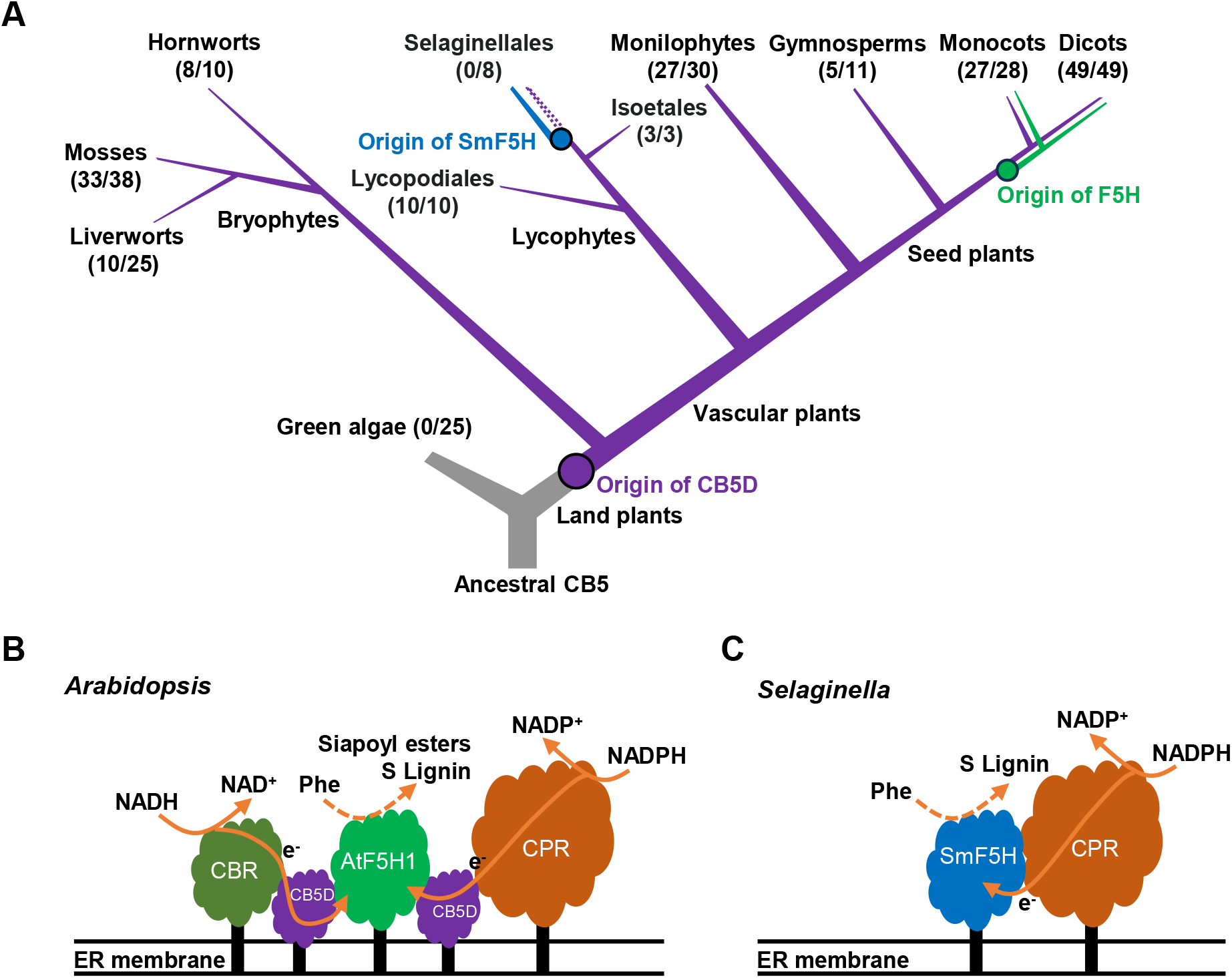
Schema of CB5D and F5H evolution and S-lignin biosynthetic cytochrome P450 systems in the basal and higher vascular plants. **A)** The evolution history of CB5D and F5H in green plants. A simplified version of the plant phylogenetic tree indicates the origin of CB5D and the phylogenetic relationship of the CB5D and F5H. The numbers in brackets indicate the conservation of H5-D/E motif in the searched species. For a given species, if it has at least one CB5 gene containing H5-D/E motif, it will be counted as one. Otherwise, this plant species will be counted as zero (see detailed information in Supplemental Data S4). **B)** and **C)** Cytochrome P450 system and the electron transfer pathway(s) for F5H-catalyzed phenolic synthesis in *Arabidopsis* **(B)** and *Selaginella* **(C)**. AtF5H1 employs both NADPH-CPR and NADH-CBR pathways, in which CB5D acts as an electron shuttle hub delivering electrons from NADPH-CPR and NADH-CBR pathways to AtF5H1 for sinapoyl esters and S-lignin biosynthesis. Whereas SmF5H only requires CPR as the sole redox partner for its catalysis.

## Discussion

### The *bona fide* CB5D evolved prior to the split of bryophyte lineages

The emergence of vascular plants on Earth occurred approximately 450 million years ago (Renault et al. 2019). A hallmark of vascular plants is the presence of the phenolic lignin heteropolymer in xylem and other sclerified cell types (Renault et al. 2019). Unlike H- and G-lignin subunits that are present in all vascular plants, S-lignin is taxonomically restricted to certain lineages, such as angiosperms and Selaginellaceae. Some species of ferns and gymnosperms have been reported to produce a trace amount of S-lignin in old biochemical studies, but this remains unverified (Weng et al. 2008b). Accordingly, the invention of F5H (CYP84A1), along with the rise of S-lignin, has been regarded as a recent evolutionary event, occurring approximately along with the emergence of angiosperm in the later of Devonian period, about 350 million years ago (Weng and Chapple 2010; Renault et al. 2019). In stark contrast to the anciently evolved cytochrome P450 C4H and C3’H in phenylpropanoid biosynthetic pathway (Renault et al. 2017; Alber et al. 2019), the more recently evolved F5H uniquely requires CB5 as an indispensable redox partner for its catalysis (Zhao et al. 2023). CB5D is a member of the CB5 family that was recruited by F5H in *Arabidopsis* and physically associated with each other for S-lignin biosynthesis (Gou et al. 2019). It is logical to postulate that this specialized CB5D redox partner might have co-evolved with F5H (CYP84A1) and emerged with the invention of F5H-catalyzed S-lignin biosynthesis. Contrary to this intuitive view, our present study reveals that the *bona fide* CB5D evolved in the ancient period of plant terrestrialization, most likely prior to the split of bryophyte lineages, therefore its occurrence predates the emergence of angiosperm F5H and S-lignin biosynthesis.

Sequence searching and phylogenetic reconstruction discover that embryophyte lineages, in contrast to their aquatic progenitor, contain at least two copies of *CB5* sequences, indicating that *CB5* gene has substantially expanded probably via gene duplication in the pioneering land plant species. Accompanied *CB5* sequence expansion is the functional divergence and specialization. Validated with genetic restoration of *Arabidopsis cb5d-1* mutant deficient in syringyl type of phenolics biosynthesis and with the augment of F5H catalytic activity in *E. coli* whole-cell biocatalytic assays (Fig. 2 and 3), the *bona fide* CB5D has already emerged in the primitive bryophytes liverwort and moss species; and was preserved in monilophytes (*S. cucullata*, *A. evecta* and *A.aleuticum*), gymnosperms (*P. glauca* and *P. abies*), and angiosperms (*Arabidopsis*, poplar, petunia and rice) (Fig. 2). Two liverwort CB5s displayed distinct subcellular localization, with one (MpCB5-1) in chloroplast, implicating its ancient origination from endosymbiotic unicell progenitor, and the another at ER membrane (MpCB5-2) that exhibits the *bona fide* CB5D function able to fully support *Arabidopsis* F5H activity and resort *cb5d* defects, indicating their functional differentiation (Fig. 2 and 3). Such functional divergence similarly occurred in moss species and was inherited in the descendant higher plants. *Arabidopsis* maintains a chloroplast localized CB5 (AtCB5A) and four ER-resident members (AtCB5B to E) (Maggio et al. 2007), including the S-lignin biosynthesis-required AtCB5D that can be functionally substituted with liverwort MpCB5-2 (Fig. 2). The basal bryophytes emerged approximately 500 million years ago (Morris et al. 2018; Renault et al. 2019). Therefore, the invention of CB5D is evolutionarily much earlier than the emergence of S-lignin biosynthetic F5H (CYP84A1). The presence of *bona fide* CB5D in both liverwort and moss species implies that the functional divergence of CB5 occurred before the bryophyte lineage split and speciation.

Bryophytes synthesize soluble phenolics such as flavonoids to deal with UV irradiation and oxidative stresses, but they lack true lignin (Basile et al. 1999; Umezawa 2003). The model moss *P. patens* contains significant amounts of caffeate-derived monomers in its cuticle (Buda et al. 2013) and a CYP98 ortholog catalyzes the phenolic ring *meta*-hydroxylation to yield caffeate derivatives in *P. patens* (Renault et al. 2017). It remains to determine the biological significance of the invention of *bona fide* CB5D in those basal land plants. CB5D in *Arabidopsis* is not required for flavonols biosynthesis (Gou et al. 2019). Nevertheless, it is interesting to examine whether or not the invented CB5D in bryophytes participates in the P450-mediated biosynthetic processes for flavonoids and/or other soluble phenolics formation. On the other hand, as a conserved electron transfer component, CB5 is well documented to associate with cytochrome *b_5_* reductase for lipid biosynthesis in yeast and mammals (Schenkman and Jansson 2003). During land colonization, the pioneering plants have to deal with desiccation challenge by inventing more complex biochemical capacities such as lipidic cuticle polymer as surface barrier to control water and gas movement. Expanding and diversifying CB5 functionalities might be to meet the biosynthetic needs for the formation of more complicated cuticle polyesters and cuticular wax, in addition to the conventional structural lipids.

### CB5D undergoes further species-specific divergence after its invention

Phylogenic analysis indicates that *CB5B/D*-like gene sequences are resolved across all the evolutionarily characteristic embryophytes, suggesting a highly conserved evolution. Nevertheless, the *bona fide* CB5D function that supports F5H catalysis for syringyl type of phenolics synthesis is not always conferred for all the CB5B/D-like proteins. This is clearly exemplified with lycophyte *Selaginella* CB5 proteins. Within well-defined *Selaginella* genomic sequences, only two *CB5* members can be identified, and *SmCB5-1* is particularly clustered within *CB5B/D*-like clade (Fig. 1). Both the encoded proteins of *SmCB5-1* and *SmCB5-2* are located to the ER membrane; however, neither of them rescued *cb5d* defects when they were expressed in *Arabidopsis* and showed function in supporting AtF5H1 or the native SmF5H activity in whole-cell biocatalytic assays (Fig. 4-6). Similarly, the aquatic floating fern *S. cucullata* has evolved four *CB5B/D*-like gene sequences. With two being tested, *S. cucullate* CB5-1 but not *S. cucullate* CB5-2 exhibited *bona fide* CB5D function (Fig. 2). These data suggest that CB5D after its early invention evolved and diverged further in the more advanced land lineages; on the other hand, its function might have been lost in some lineages such as *Selaginellales* probably due to the lack of evolutionary constraint.

### The acidic amino acid residues of helix 5 are crucial for sustaining *bona fide* CB5D function

Examining sequence deviations of CB5B/D-like proteins, it turns out that the helix 5 sequences of the CB5 proteins vary greatly in their acidic amino acid residues. The characteristic H5-D/E motif is conserved in the proteins exhibiting the *bona fide* CB5D function. The acidic amino acid residues (Asp73 and Glu74 in AtCB5D) of helix 5 significantly influence the functional specialization of CB5B/D-like proteins. The CB5B/D-like SmCB5-1 lost the corresponding Asp and Glu residues in the helix 5 region does not possess *bona fide* CB5D function; by contrast, reverse mutation of the corresponding amino acid residues to the Asp and Glu confers substantial *bona fide* CB5D functionality that substantially enhanced AtF5H1 activity and restored *cb5d* deficiency (Fig. 7). Therefore, H5-D/E motif represents a key sequence element and an evolution indicator for evolving genius CB5D protein. Notably, H5-D/E motif is ubiquitously found within nearly all land plant lineages except *Selaginellales*; and it is significantly proliferated and popularized in angiosperms (Fig. 8A), which further suggests that the *bona fide* CB5D is innovated in the early stage of land plant evolution and is largely conserved and flourished in the higher plants.

Charged acidic amino acid residues are often implicated in the intra/intermolecular electron/proton transfer processes (Kawano et al. 1998). Presumably, the distinctly positioned acidic charged residues within helix 5 might impact the redox potentials or other electrochemical or structural properties of the CB5 protein, thus defining its abilities as an electron donor for the associated oxidative enzyme. This hypothesis is seemingly supported by our *in vitro* reduction assay. The mutant variant SmCB5-1-H5 that bears Asp and Glu substitutions in its helix 5 demonstrated a more effective and steady reduction when it was treated with NADPH-dependent ATR2 (Fig. 7).

### *Selaginella* F5H does not require CB5D for its catalysis

*Selaginella moellendorffii*, a basal vascular plant, is able to synthesize S-lignin (White and Towers 1967; Weng et al. 2008b) and an independently evolved lineage-specific P450, SmF5H, functions as a dual meta-hydroxylase that complements AtF5H1 deficient mutant *fah1* (Weng et al. 2008b). Strikingly, when SmF5H were expressed in *fah1-2 cb5d-2* double mutant under the control of C4H promoter, it more effectively restored syringyl type of phenolics synthesis compared to expressing AtF5H1 driven by the same C4H promoter (Fig. 5). This functional discrepancy does not appear to be caused by their different enzymatic properties, since both F5Hs kinetically work on coniferyl alcohol and coniferaldehyde substrates equally well (Weng et al. 2010). Instead, it hints at their differences in the requirement for redox cofactors. The loss of CB5D clearly affected AtF5H1 more than SmF5H (Fig. 5). With reconstruction of P450-electron donor system in *E. coli*, AtF5H1, as our previous report, strictly required CB5D for its catalysis; while SmF5H activity was not affected at all by either AtCB5D or both SmCB5s; instead, it was strongly enhanced by NADPH-dependent CPR (Fig. 6). These lines of evidence unambiguously suggest that the independently evolved SmF5H, more like the anciently emerged C4H and C3’H and other conventional P450s, relies on NADPH-CPR electron transfer system (Fig. 8B), which differs sharply from its angiosperm counterparts F5Hs.

SmF5H has a quite dissimilar protein sequence that represents a unique *Selaginella* P450 clade (Weng et al. 2008b). Although the predicted overall structures and key structural elements, such as the E-R-R triad (in K-helix and PERF domain) and heme-binding domain (Bak et al. 2011) of AtF5H1 and SmF5H look alike, the lengths of their α-helices 8 and 10 differ substantially, let alone the large numbers of disparate amino acids (Supplemental Fig. S13). Such structural differences might alter the interfaces or the local conformations of the proteins that mediate the physical association with electron donor proteins or change redox potentials/electrochemical properties of the proteins thus influencing their recruitment on electron transfer systems. Resolving the structures of P450-redox partner complexes and further determining their electrochemical properties will shed light on understanding their unique redox catalytic mechanisms.

Taken together, our study discovers the early emergence and functional divergence of CB5D in land plants, which is sharply contrast to the recent invention of angiosperm CYP84A1 F5H. The data suggest that angiosperm F5H hijacks the anciently evolved electron transfer component to constitute a modern biochemical machinery for biosynthesis of the re-emerged S-lignin. On the other hand, the primitive F5H in the basal vascular species *Selaginella* acts as a conventional cytochrome P450 monooxygenase system that adopts the general NADPH-CPR electron transfer chain for the independently evolved S-lignin biosynthesis. These data hint that the constitution of F5H-CB5D catalytic system in angiosperm might stand an evolutionary advantage.

in this paper is available from the lead contact upon request.

## Materials and methods

### Plant materials and growth conditions

The plant materials used in this study include *N. benthamiana* and *A. thaliana.* All mutants and transgenic lines are in the WT Col-0 background. Mutant lines used were *cb5d-1* (SALK_045010), *cb5d-2* (GABI_328H06) and *fah1-2*. Transgenic lines used were listed elsewhere. Homozygous T-DNA insertion mutants were obtained by genotyping genomic DNA as a template. High order mutants were created via crossing the single mutants to obtain F1 generation seeds then the double mutants were obtained from F2 progenies via genotyping through PCR or sequencing, using the primers listed in Supplemental Data Set 1. We sterilized seeds with 70% alcohol and sowed them on half Murashige and Skoog medium (Phytotech, M524), containing 1% agar and 1% sucrose for seed germination. After 3 days of stratification at 4 °C, the seeds were germinated and maintained at 22 °C under a 16-hour light/8-hour dark regime in a growth chamber (BioChambers) with light intensity at 8500 lumens/m^2^. Seven-day-old seedlings were transferred to soil and grown until maturity under the same conditions.

### Phylogenetic analysis

All CB5 homologous sequences were retrieved by BLASTp search using *AtCB5D* (*At5g48810*) as the initial query from UCSC Genome Browser (https://genome.ucsc.edu/, *Homo sapiens*, *Mus musculus* and *Danio rerio*), Saccharomyces Genome Database (https://www.yeastgenome.org/, *Saccharomyces cerevisiae*), Phytozome 13 (https://phytozome-next.jgi.doe.gov/, *Petunia axillaris*, *Populus trichocarpa*, *Oryza sativa*, *Amborella trichopoda*, *Selaginella moellendorffii*, *Marchantia polymorpha*, *Physcomitrium patens, Volvox carteri* and *Chlamydomonas reinhardtii*), PLAZA (https://bioinformatics.psb.ugent.be/plaza/, *Picea glauca*, *Picea abies*, *Pinus taeda*, and *Ginkgo biloba*), ONEKP (https://db.cngb.org/onekp/, *Angiopteris evecta* and *Adiantum aleuticum*) and FernBase (https://www.fernbase.org/, *Salvinia cucullata*). Then the hits were reciprocally blasted (via BLASTp) against the *A. thaliana* protein sequences and discarded if the best hits were not *AtCB5A, AtCB5B, AtCB5C, AtCB5D* or *AtCB5E*. Finally, the obtained each hit was manually examined for an intact heme binding domain and the existence of a C-terminal transmembrane domain, as the criteria to be a CB5 protein. The obtained 78 CB5 coding sequences were aligned (full list in Supplemental Data Set 2) using MUSCLE algorithm integrated within the MEGA X program (Kumar et al. 2018). CB5 phylogeny tree was inferred using Maximum Likelihood method implemented in IQ-TREE-2.2.2.7 program (Minh et al. 2020; Hoang et al. 2018). Phylogenetic tree reconstruction employed the following command line: “bin\iqtree2 -s CDS-full.FAS -alrt 1000 -B 1000”. TIM2e+I+R4 model was used for the phylogenetic reconstruction which was selected by the integrated ModelFinder algorithm. 1000 ultrafast bootstrap and 1000 SHaLRT tests were performed to determine branch support (Supplemental File 1 and 2). The tree visualization and annotation were performed in TVBOT (Xie et al. 2023).

### Plant transformation

The coding sequences of *CB5* and *F5H* genes from representative plant taxa were synthesized and ligated to *pMDC32-pC4H* vector via *KpnI/SalI* restriction enzyme sites by GeneScript (https://www.genscript.com). The full list of synthesized genes is in Supplemental Data Set 3. All constructs were transformed to the corresponding homozygous *cb5d-1*, *fah1-2* or *fah1-2 cb5d-2* mutants using the agrobacterium-mediated floral dip method (Clough and Bent 1998). The primary transformants were screened on ½ MS plates containing 15 µg ml^-1^ hygromycin B (Gold Biotechnology). After 2 weeks, the positive seedlings were transferred to soil and grown to mature in a growth chamber with the above-described conditions. For the multi-gene expression of SmF5H and SmCB5s, T2A self-cleaving peptide sequence was used to fuse SmF5H and SmCB5s. The resulting DNA fragments were ligated to pMDC32-pC4H vector via *KpnI/SalI* sites. The T2A coding sequence was synthesized within the primers together with the gene specific nucleotides (Supplemental Data Set 1). SmCB5-1-H5 and SmCB5-2-H5 variants were obtained by site-directed mutagenesis using Gibson assembly method (Gibson et al. 2009).

### Soluble phenolics and lignin composition quantification

Soluble phenolics and lignin composition were quantified following the reported method (Zhao et al. 2023). Briefly, for leaf or seedling soluble phenolics analysis, about 50 mg 4-week-old rosette leaves or 1-week-old seedlings were collected and placed into a 1.5 ml centrifuge tube and weighed. 300 µl of 80% (v/v) methanol containing 80 µM chrysin (as an internal standard) and two steel balls were added. The samples were then ball-milled for 2 minutes at 30 Hz using a CryoMill (Retsch). Subsequently, an additional 200 µl of 80% (v/v) methanol containing 80 µM chrysin was added. After incubation at 4 °C for 4 hours, 5 µl supernatant was injected into a UHPLC system (Thermo Fisher Scientific) or a HPLC (Agilent) system with same settings as described (Zhao et al. 2023).

For seed soluble phenolics analysis, about 10 mg of dry seeds were mixed with 300 µl of 50% methanol containing 1.5% acetic acid and 100 µM chrysin (as an internal standard). The mixture was then subjected to ball-milling using a CryoMill (Retsch) for 2 minutes at 30 Hz, followed by the addition of 900 µl of 50% methanol containing 1.5% acetic acid and 100 µM chrysin to each sample. The samples were incubated in a cold room for 1 hour then centrifuged at 20,000g twice using an Eppendorf 5424R centrifuge. 1 µl extracts were injected into the HPLC (Agilent) system with same settings as described (Zhao et al. 2023).

For stem lignin analysis, the 10 cm basal stems from 11-week-old *Arabidopsis* plants were harvested, lyophilized and ball-milled. The obtained fine powders were extracted with 70% ethanol at 65 °C for 3 hours with three repeats, chloroform/methanol (1:1, v/v) at room temperature for 1 hour with three repeats, acetone at room temperature for overnight. The residues were dried at room temperature to obtain extractive free cell wall residues. Approximately 10 mg of extractive-free residues were weighed and placed into a glass vial, followed by addition of a freshly prepared reaction mixture composed of 2.5% boron trifluoride etherate and 10% ethanethiol in dioxane (v/v). The vials were purged with nitrogen gas before being sealed then placed in a 95 °C heat block for 4 hours with intermittent shaking. After cooling, 0.3 ml of 0.4 M sodium bicarbonate, 2 ml of water, and 1 ml of methylene chloride (containing 1 mg/ml tetracosane) were sequentially added. The vials were vortexed for 1 minute and centrifuged to separate the phases. The organic phase (approximately 1 ml) was collected and dried in a heat block overnight at 50°C. Subsequently, 0.5 ml of methylene chloride was added to resuspend the dried sample, and 50 µl of the sample was transferred to a new centrifuge tube and dried under the same condition. The sample was then derivatized by adding 50 µl of pyridine and 50 µl of N-methyl-N-(trimethylsilyl) trifluoroacetamide, followed by incubation at room temperature for 5 hours. 1 µl product was injected to an Agilent 7890A gas chromatography-flame ionization detector with same settings as described (Zhao et al. 2023).

### Subcellular localization imaging

The synthesized *CB5* genes of *M. polymorpha* and *S. moellendorffii* were amplified with primers listed in Supplemental Data Set 1 and subcloned into pDONR207 vector. After sequencing confirmation, the *CB5* genes were subcloned to pEarleyGate 104, resulting in *p35S::YFP-SmCB5-1*, *p35S::YFP-SmCB5-2*, *p35S::YFP-MpCB5-1* and *p35S::YFP-MpCB5-2* constructs. Agrobacterium strains (GV3101) carrying the constructs were transiently infiltrated to *N. benthamiana* leaves. After 3 days incubation, the fluorescence images were captured with a TCS SP5 laser-scanning confocal microscope (Leica) with excitation at 514 nm and an emission wavelength of 520 to 535 nm for YFP signals. The chloroplast autofluorescence was observed at 636 to 725 nm.

### RT-qPCR analysis of gene expression

Total RNAs were extracted from plant materials in triplicate using TRIzol^TM^ reagent (ThermoFisher Scientific) following the manufacturer’s instruction. Reverse transcription reaction was conducted using 0.5 µg of total RNA and 2 µl of iScript™ Reverse Transcription Supermix (Bio-Rad) in 10 µl reaction volume. The reaction mixture was incubated in a thermal cycler at 25 °C for 5 min, 46 °C for 20 min, and 95 °C for 1 min. The cDNA solution was diluted 5 times, and 2 µl of the diluted solution was used as template in a 15 µl reaction with SsoAdvanced Universal SYBR Green Supermix (Bio-Rad). qPCR was performed using CFX96 Real-Time System (Bio-Rad) and cycle threshold value was calculated by the CFX Manager Software v.3.3 (Bio-Rad). Primers used in qPCR were listed in Supplemental Data Set 1. *Arabidopsis PP2A* gene was used as the housekeeping reference gene and the data were calculated using the delta-cycle or delta-delta-cycle threshold method (Schmittgen and Livak 2008).

### Protein expression, purification, and reduction assay

The expression and purification of AtCBR1 and ATR2 were described as previous (Zhao et al. 2023). For SmCB5s and their variant and MpCB5s, the coding sequences with removal of transmembrane domain were PCR-amplified and inserted to pET28a (+) vector via *EcoRI* and *XhoI* restriction enzyme sites. The protein expression and purification of CB5 proteins and the reduction assay were performed as described previously (Zhao et al. 2023). Briefly, the sequence verified constructs were transformed to *E. coli* BL21 cells, inoculated to 20 ml Terrific Broth and cultured overnight at 37 °C. The precultures were inoculated to 600 ml Terrific Broth and grew at 37 °C to an optical density of around 1 at 600 nm. After 30 min culture at 15 °C, 0.5 mM isopropyl-β-D-thiogalactopyranoside (IPTG) was added for protein induction. Overnight cultures were harvested and stored at −80 °C freezer. Recombinant proteins were purified with Ni^2+^-nitrilotriacetic acid agarose beads (Qiagen) and desalted with Bio-Gel P-6DG gel (Bio-Rad). CB5 protein concentration was quantified from the different spectra of the cytochrome catalyzed by dithionite using the extinction coefficient of Δε (reduced-oxidized)_424-409_ = 185 mM^-1^·cm^-1^ or from the absolute spectrum using extinction coefficient ε_413_ = 117 mM^-1^·cm^-1^ (Naito et al. 1998). The purified protein was aliquoted and stored at −80 °C for further use. The redox assay was performed in 100 µl 20 mM Tris-HCl buffer (pH7.5) in 96-well microplate. The oxidized and reduced status of CB5 proteins were recorded before and after adding NADH or NADPH, respectively.

### Whole-cell biocatalytic assay in *E. coli*

AtF5H1 and SmF5H coding sequences with removal of their transmembrane domain were PCR-amplified and inserted to the reported pET28a(+)-CB5D and -ATR2 vector (Zhao et al. 2023) via *Bam HI* restriction enzyme site to obtain chimeric constructs, *AtF5H1-CB5D*, *SmF5H-CB5D* and *SmF5H-ATR2*. The *SmF5H* fragment was PCR-amplified and inserted to pET28a(+) vector via *EcoRI* and *XhoI* sites. The *AtF5H1*, *2muCB5D*, *SmF5H*, *SmCB5*, *SmCB5-H5* and *SmCPR* fragments were PCR-amplified using overlapping primers and inserted to pET28a(+) vector via *BamHI* and *XhoI* sites to obtain chimeric constructs, *AtF5H1-2muCB5D*, *AtF5H1-SmCB5-1, AtF5H1-SmCB5-2, AtF5H1-SmCB5-1-H5, AtF5H1-SmCB5-2-H5, SmF5H-SmCB5-1, SmF5H-SmCB5-2, SmF5H-SmCB5-1-H5, SmF5H-SmCB5-2-H5* and *SmF5H-SmCPR*. The green algal *CrCB5* and *VcCB5* genes (Supplemental Data Set 3) were synthesized and ligated to the pET28a(+)-AtF5H1-CB5D vector (GeneScript) by replacing *CB5D* fragment via *EcoRI* and *XhoI* sites to obtain chimeric constructs *AtF5H1-CrCB5* and *AtF5H1-VcCB5*. The obtained constructs were transformed into *E. coli* BL21 cells. 500 µl of overnight cultures were inoculated to 10 ml terrific broth and grew at 37 °C until an optical density of ∼1 at 600 nm. The cells grew at 15 °C for 30 min before induction with 1 mM IPTG. Overnight cultures were pelleted and resuspended with 3 ml of potassium phosphate buffer (50 mM; pH 7.4) containing 2% glucose. 60 nmol coniferyl alcohol was added as F5H enzyme substrate. The samples were incubated at 28 °C with 250 rpm rotation for 4 hours. The reactions were stopped by adding 1 ml of ethyl acetate containing 10 µM chrysin as the internal standard. The extracts were vacuum-dried and analyzed with an Agilent 1100 HPLC system with the same settings as described above.

### Statistical analysis

Statistical analysis in each required experiment was performed using either Student’s t-test with Microsoft Excel (two-tailed distribution and two-sample unequal variance) or analysis of variance (ANOVA) test with GraphPad Prism version 4 (*P* < 0.05, one-way ANOVA and Tukey’s test). Details of statistical analyses, including sample sizes and biological replications, are provided in the figure legends. Detailed statistical analysis data are shown in Supplemental Data Set 5.

## Accession numbers

DNA and derived protein sequence data from this article are available in the UCSC Genome Browser (https://genome.ucsc.edu/), Saccharomyces Genome Database (Saccharomyces cerevisiae), TAIR database (https://www.arabidopsis.org/), Genebank (https://www.ncbi.nlm.nih.gov/genbank/), Phytozome 13 (https://phytozome-next.jgi.doe.gov/), PLAZA (https://bioinformatics.psb.ugent.be/plaza/), ONEKP (https://db.cngb.org/onekp/), Solanaceae Genomics Network (https://solgenomics.net/) and FernBase (https://www.fernbase.org/). The *Arabidopsis* genes are under the following TAIR accession numbers: AtCB5A (At1g26340); AtCB5B (AT2G32720); AtCB5C (At2g46650); AtCB5D (At5g48810); AtCB5E (At5g53560); AtF5H1 (At4g36220).

## Author Contributions

C-J.L. conceived the study. C-J L. and X. Z. designed the experiments. X.Z conducted experiments. Y.Z. initiated *in planta* complementation assays. Q-Y.Z. participated in phylogenetic analysis. C-J.L. and X.Z. analyzed and interpreted data and wrote the manuscript. All authors edited the manuscript.

## Supplemental data

The following materials are available in the online version of this article.

**Supplemental Figure S1.** The simplified phenylpropanoid-monolignol biosynthetic pathway leading to the synthesis of S-lignin and sinapate esters in angiosperms and the lycophyte *Selaginella*.

**Supplemental Figure S2.** PCR and/or RT-qPCR confirmation of the *CB5* transgenic lines in *Arabidopsis cb5d-1* background.

**Supplemental Figure S3.** SDS-PAGE gel images showing the induced production of F5H-CB5 fusion proteins as labeled.

**Supplemental Figure S4.** UHPLC-MS analysis of *E. coli* whole-cell biocatalytic assays for AtF5H1 catalytic activity.

**Supplemental Figure S5.** Whole-cell biocatalytic assays for AtF5H1-CB5 fusion proteins.

**Supplemental Figure S6.** Lignin composition analysis in the representative fern and gymnosperm species and complementation assay of *fah1-2* mutant with the selected *F5H* candidates.

**Supplemental Figure S7.** Verification of *SmCB5* transgenic lines in *cb5d-1* background.

**Supplemental Figure S8.** Complementation of *fah1-2* mutant with *SmF5H*.

**Supplemental Figure S9.** Creation and confirmation of *fah1-2 cb5d-2* double mutant.

**Supplemental Figure S10.** Gene expression in the putative complementation lines of *fah1-2 cb5d-2*.

**Supplemental Figure S11.** Homology models of AtCB5D and SmCB5-1 and multisequence alignment of the genetically confirmed CB5s.

**Supplemental Figure S12.** Verification of transgenic lines harboring *SmCB5* variants in *cb5d-1* background.

**Supplemental Figure S13.** Comparison of S-lignin biosynthetic F5H enzymes of *Arabidopsis* and *Selaginella*.

**Supplemental Data Set 1.** Primers used in this study.

**Supplemental Data Set 2.** Genes used for phylogenetic analysis in Figure 1B.

**Supplemental Data Set 3.** Synthesized genes in this study.

**Supplemental Data Set 4.** The helix 5 sequences in searched species used for calculating helix 5 conservation in Figure 8A.

**Supplemental Data Set 5.** Summary of statistical analyses.

**Supplemental File 1.** Multiple sequence alignment for Figure 1B.

**Supplemental File 2.** Newick format of the phylogenetic tree for Figure 1B.

